## Supplementary Information for "Robust and tunable signal processing in mammalian cells via engineered covalent modification cycles"

Jones et al.

### 1 Supplementary Note 1: Model of TCS phosphoregulation

#### 1.1 Model of the covalent modification cycle

We consider the covalent modification cycle (CMC) shown in Figure 3a. An autophosphorylated HK (K) phosphorylates the RR (X) to become $\mathrm{X}^*$, which can transcriptionally regulate downstream genes. Conversely, a phosphatase P can de-phosphorylate $\mathrm{X}^*$ to become X. These biomolecular reactions are described by the following reactions:

$$\mathrm{X} + \mathrm{K} \underset{\mathrm{d_k}}{\overset{\mathrm{a_k}}{\rightleftharpoons}} \mathrm{c_k} \xrightarrow{\theta_\mathrm{k}} \mathrm{X}^* + \mathrm{K}, \qquad \mathrm{X}^* + \mathrm{P} \underset{\mathrm{d_p}}{\overset{\mathrm{a_p}}{\rightleftharpoons}} \mathrm{c_p} \xrightarrow{\theta_\mathrm{p}} \mathrm{X} + \mathrm{P}, \tag{9}$$

where $\mathrm{c_k}$ and $\mathrm{c_p}$ are complexes formed by the HK and the phosphatase with the RR, respectively, and $\theta_k$ and $\theta_p$ are the catalytic rate constants for the HK and the phosphatase, respectively. The RR (X) is expressed constitutively and all species decay with rate constant $\gamma$:

$$\varnothing \xrightarrow{\alpha_\mathrm{x}} \mathrm{X}, \qquad \mathrm{X}, \mathrm{X}^*, \mathrm{c_p}, \mathrm{c_k} \xrightarrow{\gamma} \varnothing. \tag{10}$$

The lumped parameter $\alpha_x$ is the production rate constant of RR that increases with its DNA copy numbers, promoter strength, and translation rate constant. The parameter $\gamma$ is the dilution rate constant. Additionally, the total concentrations of RR, HK, and phosphatase are conserved:

$$K_t = K + c_k, \qquad P_t = P + c_p, \qquad X_t = X + X^* + c_k + c_p. \tag{11}$$

A mass-action kinetics model[93] corresponding to (9)-(11) takes the following form:

$$\frac{\mathrm{d}}{\mathrm{d}t} c_k = a_k X K - d_k c_k - \theta_k c_k - \gamma c_k, \tag{12a}$$

$$\frac{\mathrm{d}}{\mathrm{d}t} c_p = a_p X^* P - d_p c_p - \theta_p c_p - \gamma c_p, \tag{12b}$$

$$\frac{\mathrm{d}}{\mathrm{d}t} X_t = \alpha_x - \gamma X_t, \tag{12c}$$

$$\frac{\mathrm{d}}{\mathrm{d}t} X^* = \theta_k c_k - a_p X^* P + d_p c_p - \gamma X^*. \tag{12d}$$

Setting the time derivatives in (12a)-(12b) to 0, the quasi-steady state (QSS) concentrations of the complexes are found to be:

$$c_k = \frac{XK}{K_{M,k}}, \qquad c_p = \frac{X^* P}{K_{M,p}}, \tag{13}$$

where the Michaelis-Menten constants are defined as:

$$K_{M,k} := \frac{d_k + \theta_k + \gamma}{a_k} \approx \frac{d_k + \theta_k}{a_k}, \qquad K_{M,p} := \frac{d_p + \theta_p + \gamma}{a_p} \approx \frac{d_p + \theta_p}{a_p}.$$

The above approximations are taken assuming that the catalytic rate constants are much faster than the decay rate constants: $\theta_k, \theta_p \gg \gamma$. Given (11) and (13), the concentrations of free kinase and phosphatase can be computed as:

$$K = K_t \cdot \frac{K_{M,k}}{K_{M,k} + X}, \qquad P = P_t \cdot \frac{K_{M,p}}{K_{M,p} + X^*}. \tag{14}$$

If we assume that the substrates are much more abundant than the enzymes so that free enzymes are limited: $K \ll K_{M,k}$ and $P \ll K_{M,P}$, then $c_k \ll X$ and $c_p \ll X^*$ so that (11) becomes:

$$X_t = X + X^*. \tag{15}$$

Substituting (13) and (14) into (12d), we derive the dynamics of $X^*$:

$$\frac{\mathrm{d}}{\mathrm{d}t}X^* = \theta_k \frac{XK}{K_{M,k}} - \theta_p \frac{X^*P}{K_{M,p}} - \gamma X^* = \theta_k \frac{(X_t - X^*)K_t}{(X_t - X^*) + K_{M,k}} - \theta_p \frac{X^* P_t}{X^* + K_{M,p}} - \gamma X^*. \tag{16}$$

We model transcription activation of the output protein Y by X* through a Hill function[93]. Considering decay of the output protein, its dynamics follow:

$$\frac{\mathrm{d}}{\mathrm{d}t}Y = \alpha \frac{(X^*/K)^n}{1 + (X^*/K)^n} - \gamma Y, \tag{17}$$

where $\alpha$ is the maximum production rate constant of the output, $K$ is the dissociation constant describing binding between X* and the promoter driving the output, and $n$ is the Hill cooperativity.

#### 1.2 Model of the disturbance inputs

For simplicity, we model disturbances created by post-transcriptional repression (created by miR-FF4) and by resource loading (created by Gal4-VPR expression) as a fold change ($w \in [0, 1)$) in the protein production rates. In particular, if the output mRNA is targeted by miR-FF4, the production rate of the output gene only is affected by the disturbance. Hence, we write the open loop system model as:

$$\frac{\mathrm{d}}{\mathrm{d}t}X_t = \alpha_x - \gamma X_t, \quad \frac{\mathrm{d}}{\mathrm{d}t}X^* = \theta_k \frac{(X_t - X^*)K_t}{(X_t - X^*) + K_{M,k}} - \theta_p \frac{X^* P_t}{X^* + K_{M,p}} - \gamma X^*, \quad \frac{\mathrm{d}}{\mathrm{d}t}Y = \alpha(1 - w)\frac{(X^*/K)^n}{1 + (X^*/K)^n} - \gamma Y. \tag{18}$$

On the other hand, if the disturbance is created by resource loading, the production rates of RR (X) and output (Y) are both affected. In this case, we write the open loop system model as:

$$\frac{\mathrm{d}}{\mathrm{d}t}X_t = \alpha_x(1 - w) - \gamma X_t, \quad \frac{\mathrm{d}}{\mathrm{d}t}X^* = \theta_k \frac{(X_t - X^*)K_t}{(X_t - X^*) + K_{M,k}} - \theta_p \frac{X^* P_t}{X^* + K_{M,p}} - \gamma X^*, \quad \frac{\mathrm{d}}{\mathrm{d}t}Y = \alpha(1 - w)\frac{(X^*/K)^n}{1 + (X^*/K)^n} - \gamma Y. \tag{19}$$

#### 1.3 Model of the phosphatase feedback system

When the CMC is in a feedback configuration, with the phosphatase production regulated by the active RR (X*), the total amount of phosphatase ($P_t$) is no longer a constant, but instead dependent on the concentration of the active

RR ($X^*$). In particular, the dynamics of the total amount of phosphatase follow:

$$\frac{\mathrm{d}}{\mathrm{d}t}P_t = \alpha_p \frac{(X^*/K)^n}{1+(X^*/K)^n} - \gamma_p P_t, \tag{20}$$

where $\alpha_p$ is the production rate constant of the phosphatase and $\gamma_p$ is the phosphatase decay rate constant that can be modulated by TMP. Specifically, the magnitude of $\gamma_p$ decreases with increasing TMP.

When the closed loop system is subject to a disturbance created by miR-FF4, its dynamics follow:

$$\begin{aligned}
\frac{\mathrm{d}}{\mathrm{d}t}X_t &= \alpha_x - \gamma X_t,\\
\frac{\mathrm{d}}{\mathrm{d}t}X^* &= \theta_k \frac{(X_t - X^*)K_t}{(X_t - X^*) + K_{M,k}} - \theta_p \frac{X^* P_t}{X^* + K_{M,p}} - \gamma X^*,\\
\frac{\mathrm{d}}{\mathrm{d}t}P_t &= \alpha_p(1-w)\frac{(X^*/K)^n}{1+(X^*/K)^n} - \gamma_p P_t,\\
\frac{\mathrm{d}}{\mathrm{d}t}Y &= \alpha(1-w)\frac{(X^*/K)^n}{1+(X^*/K)^n} - \gamma Y.
\end{aligned} \tag{21}$$

When the closed loop system is subject to a disturbance created by squelching, its dynamics follow:

$$\begin{aligned}
\frac{\mathrm{d}}{\mathrm{d}t}X_t &= \alpha_x(1-w) - \gamma X_t,\\
\frac{\mathrm{d}}{\mathrm{d}t}X^* &= \theta_k \frac{(X_t - X^*)K_t}{(X_t - X^*) + K_{M,k}} - \theta_p \frac{X^* P_t}{X^* + K_{M,p}} - \gamma X^*,\\
\frac{\mathrm{d}}{\mathrm{d}t}P_t &= \alpha_p(1-w)\frac{(X^*/K)^n}{1+(X^*/K)^n} - \gamma_p P_t,\\
\frac{\mathrm{d}}{\mathrm{d}t}Y &= \alpha(1-w)\frac{(X^*/K)^n}{1+(X^*/K)^n} - \gamma Y.
\end{aligned} \tag{22}$$

### 2 Supplementary Note 2: Approaches to improve the robustness of the CL system

In our experiments, we found that our closed loop (CL) system had improved robustness to perturbations compared to comparable open loop (OL) systems, but this robustness was not perfect (Figure 6 & Supplementary Figures 23-32. In theory, the phosphatase feedback topology can achieve near-perfect adaptation via quasi-integral feedback control[27]. Perfect adaptation is achieved by having a large concentration of the phospho-TF (OmpR-VP64 in our case) and a small $K_M$ for the phosphatase relative to the $K_D$ of TF-DNA binding[27] (see Model Box). Below, we discuss the benefits and drawbacks of different approaches to realize integral feedback control with proteins derived from bacterial TCS.

#### 2.1 Increase the total amount of RR

A key condition for achieving integral feedback control with in our device is to saturate both the kinase and phosphatase[27]. The kinase can be trivially saturated by introducing a large amount of phosphoprotein ($X$). However, saturation of the phosphatase requires both a large amount of $X$ and a sufficient amount of kinase to phosphorylate $X$ to a quantity of phospho-$X$ ($X^*$) such that the phosphatase is also saturated. However, if the total amount of $X$ ($X_{Tot}$) is too low, then there will be no regime whereby both enzymes are saturated. Thus, one potential problem with our circuit implementation is that the level of OmpR-VP64 is too low, and increasing it would improve robustness. However, in the real system, the level of OmpR-VP64 cannot be too high because the non-phosphorylated protein retains ~20-fold lower affinity for the promoter[50] (Supplementary Figure 17). Thus, as OmpR-VP64 levels increase, the output level becomes decoupled from the phosphorylation state of OmpR-VP64. Other TCS systems in which the RR has a larger difference between the $K_D$ of RR and RR-P could be identified and used to avoid this issue.

#### 2.2 Increase the ratio of $K_M : K_D$

For perfect adaptation, the level of $X^*$ must be much higher than the $K_M$ of the phosphatase, yet below the $K_D$ for promoter saturation[27] (see Model Box). Given that our system response to perturbation is partially but incompletely adaptive, it is likely that these values are relatively close to one another, rather than the $K_M$ being much smaller than $K_D$ as would be ideal.

Thus, one option to improve performance is to increase the $K_D$ of binding between OmpR-VP64 and DNA. However, this increases the level of OmpR-VP64 needed for activation, and high expression of transcriptional activators can lead to gene expression burden on the cell and a reduction in output capacity due to cellular resource loading[51,63,94–96]. In addition, producing enough OmpR-VP64 to achieve significant promoter activation can itself be challenging, given that TFs are typically unstable[97].

In addition, we could decrease the $K_M$ of the phosphatase. Mathematically, $K_M = \frac{k_{cat}+k_{off}}{k_{on}}$. The $k_{cat}$ of EnvZ kinase activity is $\sim 0.1\text{min}^{-1}$ and is limited by the autokinase rate of EnvZ[68,98]. the $k_{cat}$ of EnvZ phosphatase activity is estimated to be roughly 10 times higher (~1 $\text{min}^{-1}$)[36,99]. Compared to both $k_{cat}$ values, the off-rate for Envz-OmpR binding is estimated to be larger; the $K_D$ of EnvZ-OmpR binding is $\sim 1\mu$M[100], so assuming that the $k_{on}$ is typical for an enzyme ($\sim 10^5 M^{-1}s^{-1}$), then $k_{off} \sim 6\text{min}^{-1}$ If $k_{on}$ is higher, then in fact the $k_{off}$ would be much larger than either $k_{cat}$. Consequently, the most effective way to reduce the $K_M$ would be to reduce the $k_{off}$ by increasing the affinity of EnvZ variants for OmpR. This has been done previously using scaffolding proteins[101], and could be extended to higher-affinity interactions such as scFvs to epitope tags[102]. However, increasing the binding affinity between EnvZ and OmpR may have negative consequences due to indiscriminate binding between EnvZ and either OmpR and P-OmpR[100], which enables both the phosphatase and kinase variants of EnvZ to bind the products of their catalysis reactions. Binding of the enzymes to their products could lead to sequestration of OmpR molecules by the enzymes, and thus reduce the signaling capacity of the system by reducing the free concentrations of all three species. Thus, we would ideally increase affinity for the substrate only, and not the product. This may be possible via mutagenesis of the binding interfaces between EnvZ and OmpR, though it is unclear if it is possible to dramatically alter the binding preferences of EnvZ for different phospho-forms of OmpR.

#### 2.3 Create a phosphorylation cascade

Functionally, a high concentration of phosphoprotein compared to the $K_M$ of the enzymes leads to ultrasensitivity in the phosphorylation level to the amount of either enzyme[23]. This ultrasensitivity, in turn, provides the digital ("zero-order")[23] on-off response needed for quasi-integral feedback control. Ultrasensitivity can be alternatively achieved by constructing a cascade of multiple phosphorylation cycles[23]. One possibility for implementing such a cascade involves using hybrid RRs, which comprise both RR rec domains and HK DHp domains for phosphorelay[15,92]. However, because the phosphate groups are transferred among protein domains rather than used to covalently activate subsequent catalytic domains, this method of phosphotransfer does not actually implement the cascade as desired. Thus, to realize this design, future work will be needed to identify a novel method of inserting additional CMCs into the circuit.

### Supplementary Figures & Tables

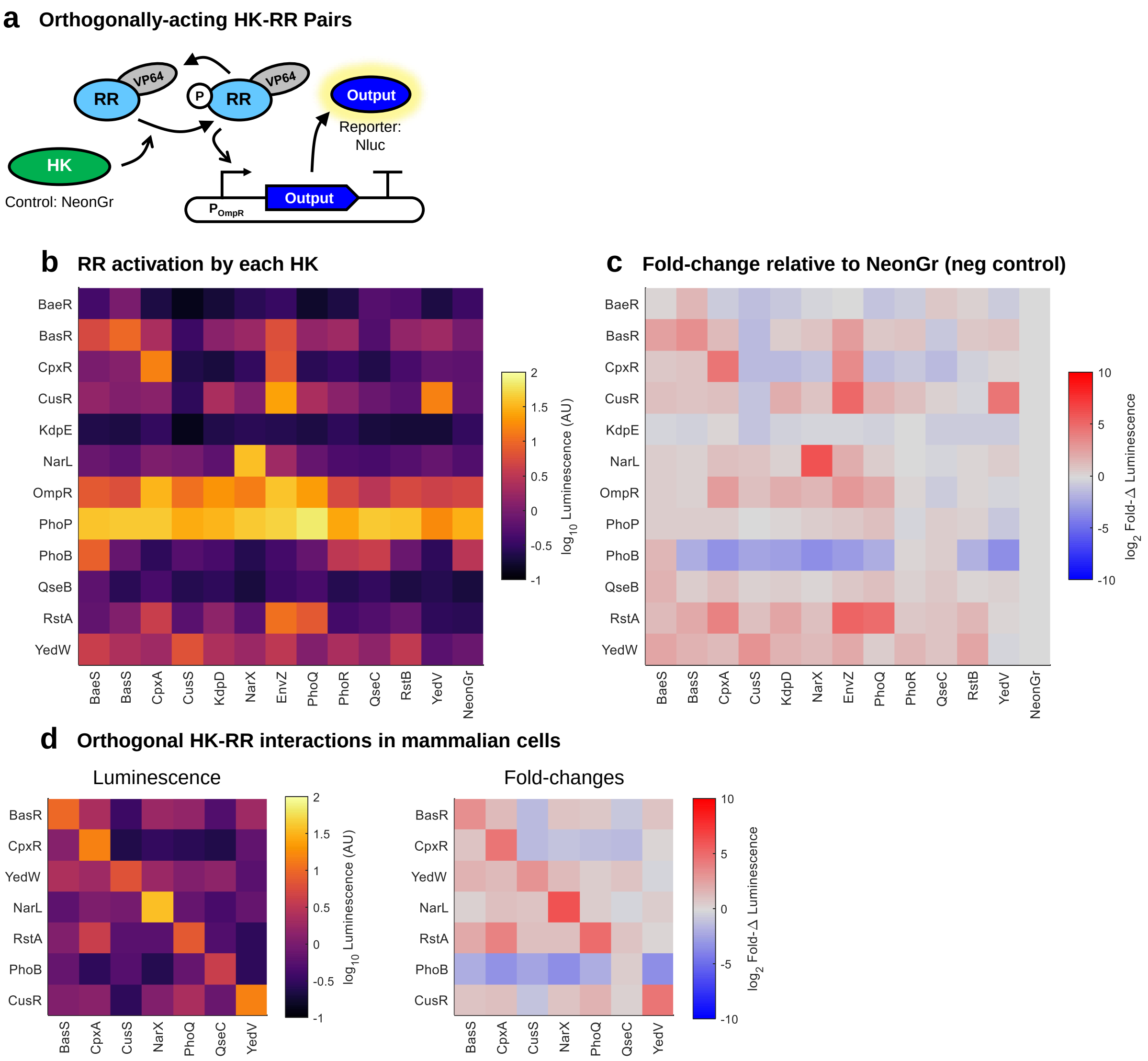


**Supplementary Figure 1: Orthogonality of TCS pathways in human cells.**

**Supplementary Figure 1:** (Previous page.) **(a)** Schematic of experiment design. Different *E. coli* TCS proteins were co-transfected with luminescent reporter plasmids bearing 6x RR binding sites upstream of a synthetic minimal promoter (minKB, *i.e.* YB_TATA)[22]. **(b)** Luminescence RR-driven promoters for each combination of HK and RR proteins. RRs were always paired with reporters driven by their cognate promoters. Luminescence was normalized by dividing by the signal from a constitutive transfection marker (hPGK:Fluc2). **(c)** Fold-changes to a control sample (mNeonGreen expressed in place of a HK). Since mNeonGreen is a fluorescent reporter and luminescence was measured, it does not contribute to the signal output. **(d)** A subset of seven HK-RR pairs with the best orthogonality among all HK-RR combinations. Luminescence values are shown on the left and fold-changes compared to the mNeonGreen control on the right. Note that the QseC-PhoB combination appears to prevent promoter knockdown rather than activating the promoter. The PhoB promoter appears to effectively be constitutive and does not drive increased output in the presence of PhoR. The OmpR and PhoP promoters also have high basal activities, but still respond strongly to their respective inputs. Two HKs (YedV/CusS) appear to activate each other's cognate RR but not their own. Cross-talk between these pathways has been observed before[68], but the lack of self-activation is unexpected. It is possible that these HKs take on a kinase-dominant conformation when binding to the non-cognate RR and a phosphatase-dominant conformation when binding to the cognate RR. Additional cross-talk between these pathways has been seen at the DNA level – YedW and CusR co-regulate many target genes by binding nearly identical DNA sequences[103]. PhoQ appears to activate RstA, but measurement with a fluorescent reporter showed no response (Supplementary Figure 2). All data was collected 48 hours post-transfection in HEK-293FT cells.

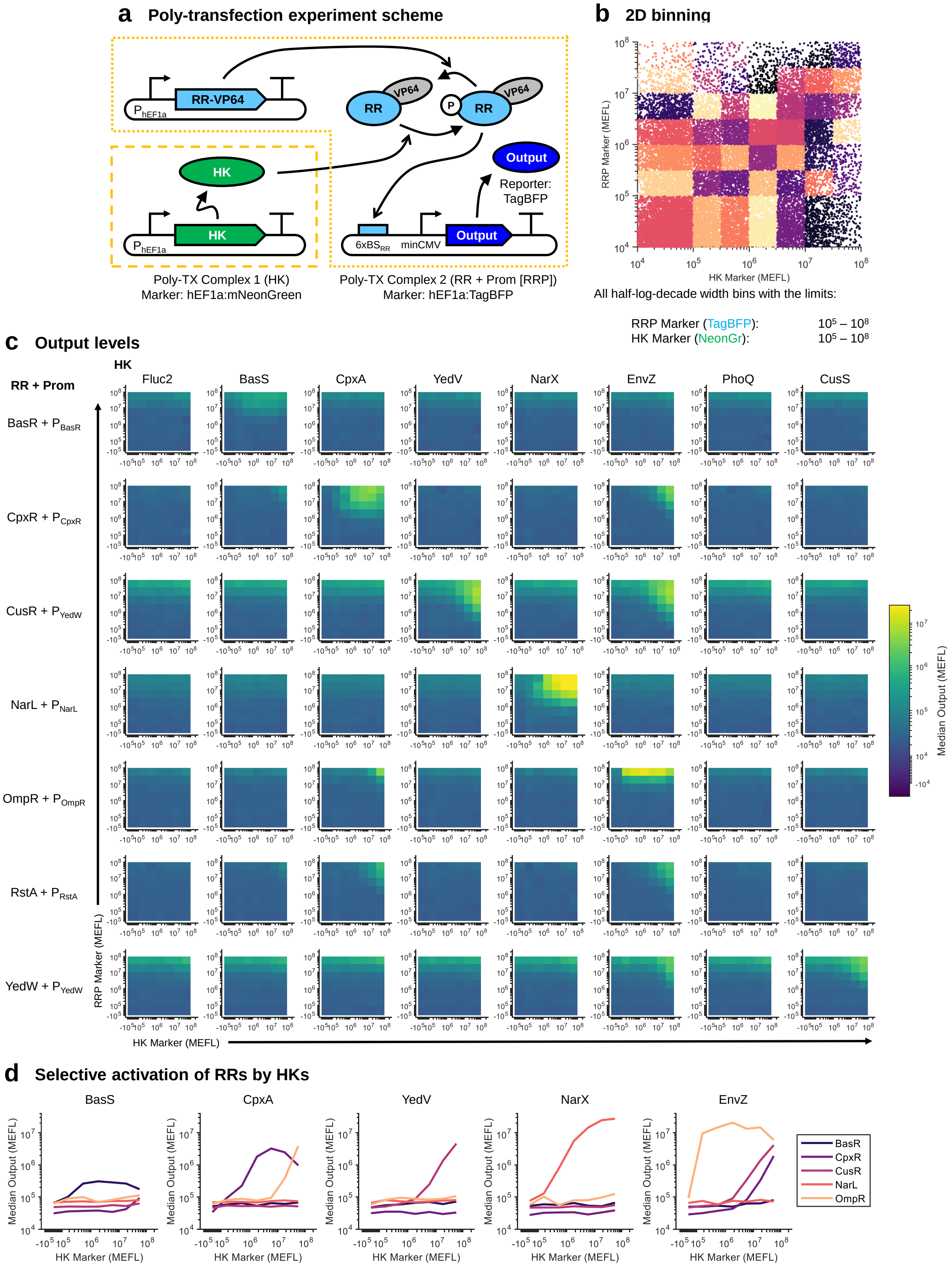


**Supplementary Figure 2: Dose responses of RRs and their cognate promoters to HK inputs**.

**Supplementary Figure 2:** (Previous page.) **(a)** Schematic for poly-transfection experiment. Different HKs and RRs found to operate orthogonally (Supplementary Figure 1d) were delivered in separate DNA-lipid complexes, thereby decoupling their delivery to cells[44] and enabling the calculation of dose-responses between the amount of HK and the level of RR-driven output. The RR plasmids and RR-activated promoters driving the output reporter were co-delivered in Complex 2, which is also referred to as "RRP". **(b)** 2D binning scheme for analysis of the poly-transfection data. The decoupled delivery of the HKs (as indicated by the HK Marker) and the other plasmids (as indicated by the RRP Marker) yield cells with various combinations of DNA dosages of each. The binning groups cells together based on DNA dosages delivered by each complex. **(c)** Median output expression level per bin for each combination tested. **(d)** Comparison of dose-responses of RR-driven outputs to each HK (omitting CusS/YedW, for which the target promoter is shared with and less strongly activated than by YedV/CusS, and PhoQ/RstA, for which no output activation was observed). The RRP Marker bin was selected uniquely for each RR/promoter to optimize on-target activation and minimize off-target activation. This same bin is used for all subplots. BasR: Bin 6 ($\approx 10^{7.25}$ MEFLs), CpxR: Bin 6, CusR: Bin 5 ($\approx 10^{6.75}$ MEFLs), NarL: Bin 6, OmpR: Bin 7 ($\approx 10^{7.75}$ MEFLs). All data was collected 48 hours post-transfection in HEK-293FT cells.

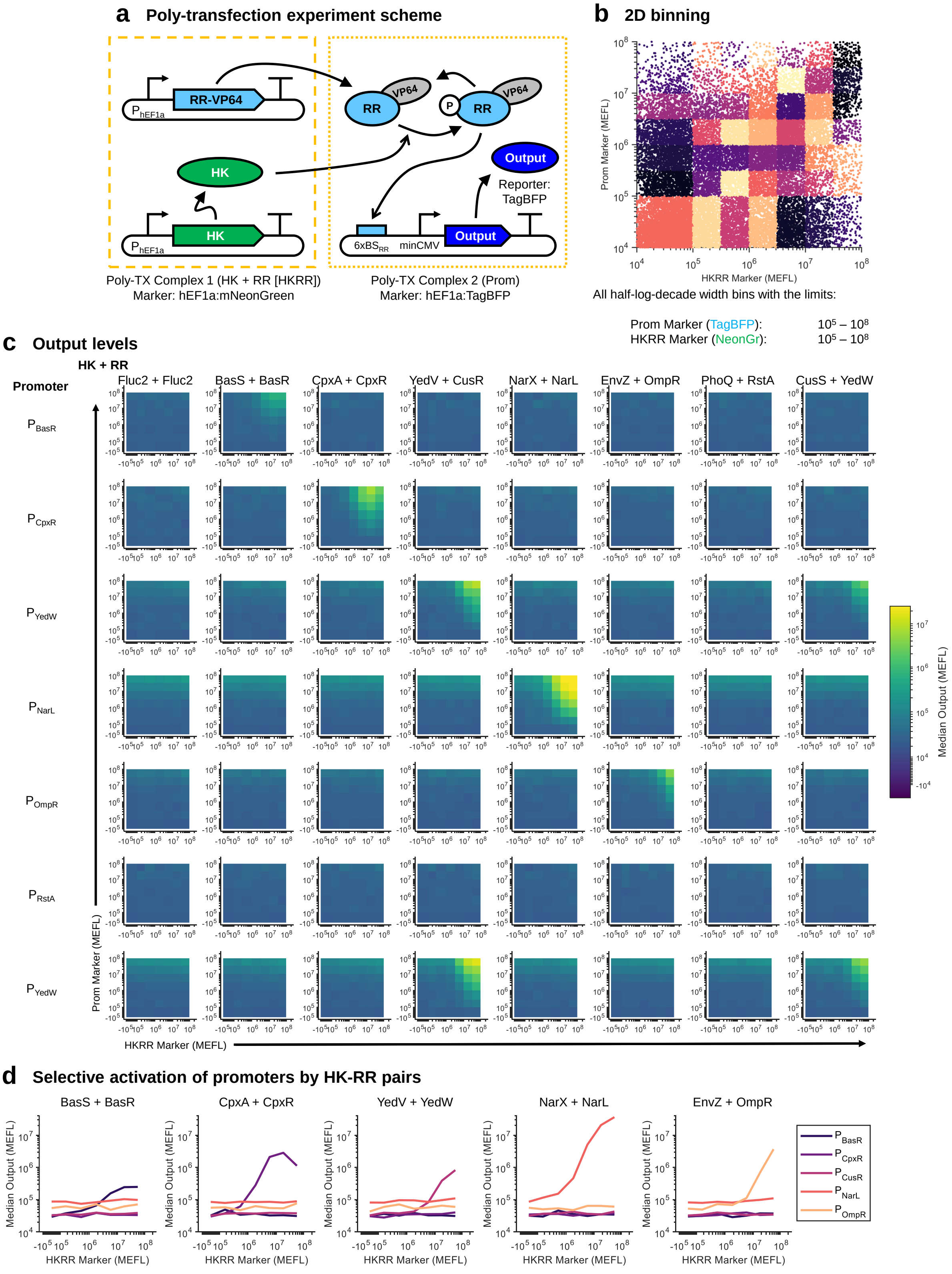


**Supplementary Figure 3: Dose responses of RR-driven promoters to HK-RR pairs.**

**Supplementary Figure 3:** (Previous page.) **(a)** Schematic of poly-transfection experiment. Similar to the experiment in Supplementary Figure 2, except with HK and RR pairs co-delivered in one complex and the RR-driven promoters delivered separately. **(b)** 2D binning scheme for analysis of the poly-transfection data. **(c)** Median output expression level per bin for each combination tested. **(d)** Comparison of dose-responses of RR-driven outputs to each HK/RR pair; the bins selected for comparison are the same as in Supplementary Figure 2d. All data was collected 48 hours post-transfection in HEK-293FT cells.

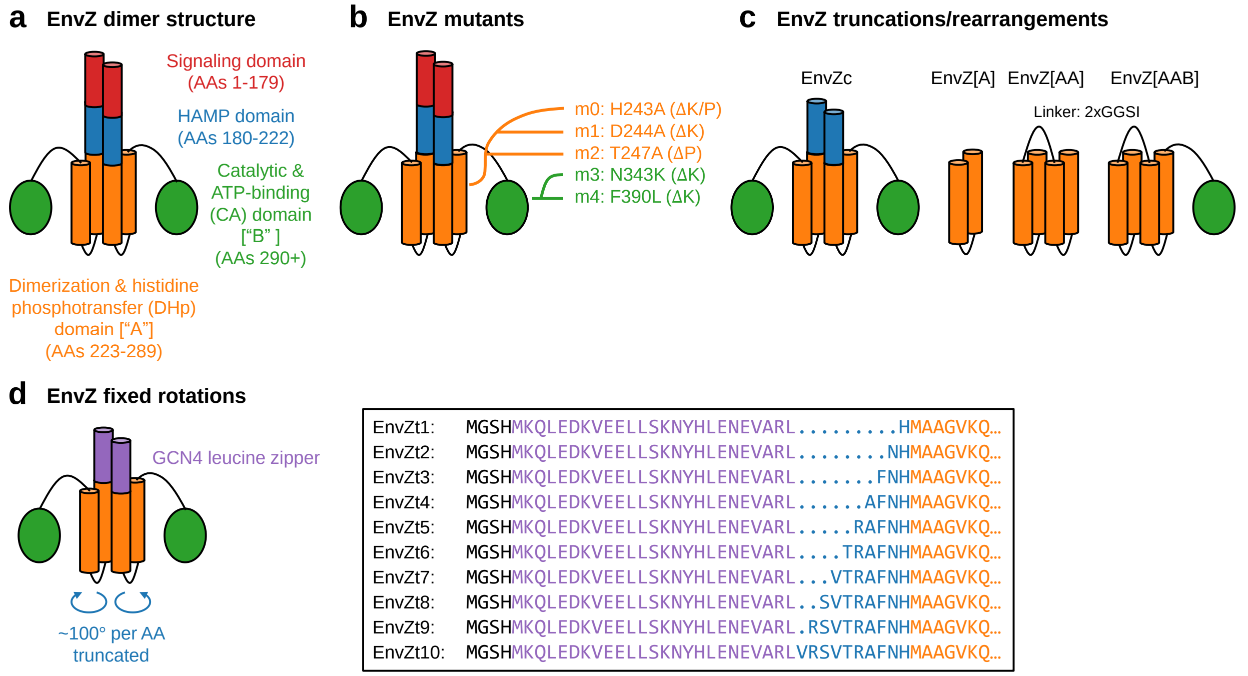


**Supplementary Figure 4: Variants of EnvZ**. **(a)** High-level structure of EnvZ, with annotation of important domains for intracellular signal transduction. **(b)** Mutants of EnvZ[32–34,39,104] tested in this study, mapped onto their location in the EnvZ structure. **(c)** Truncations[35,36] and domain rearrangements[37] of EnvZ tested in this study. **(d)** Schematic for the GCN4-rotationally-locked[40,105] variants of EnvZ; the box on the right shows the sequence of the first 37-46 amino acids of each variant. Throughout this study, we combine some of these variants together as indicated in the figures/text.

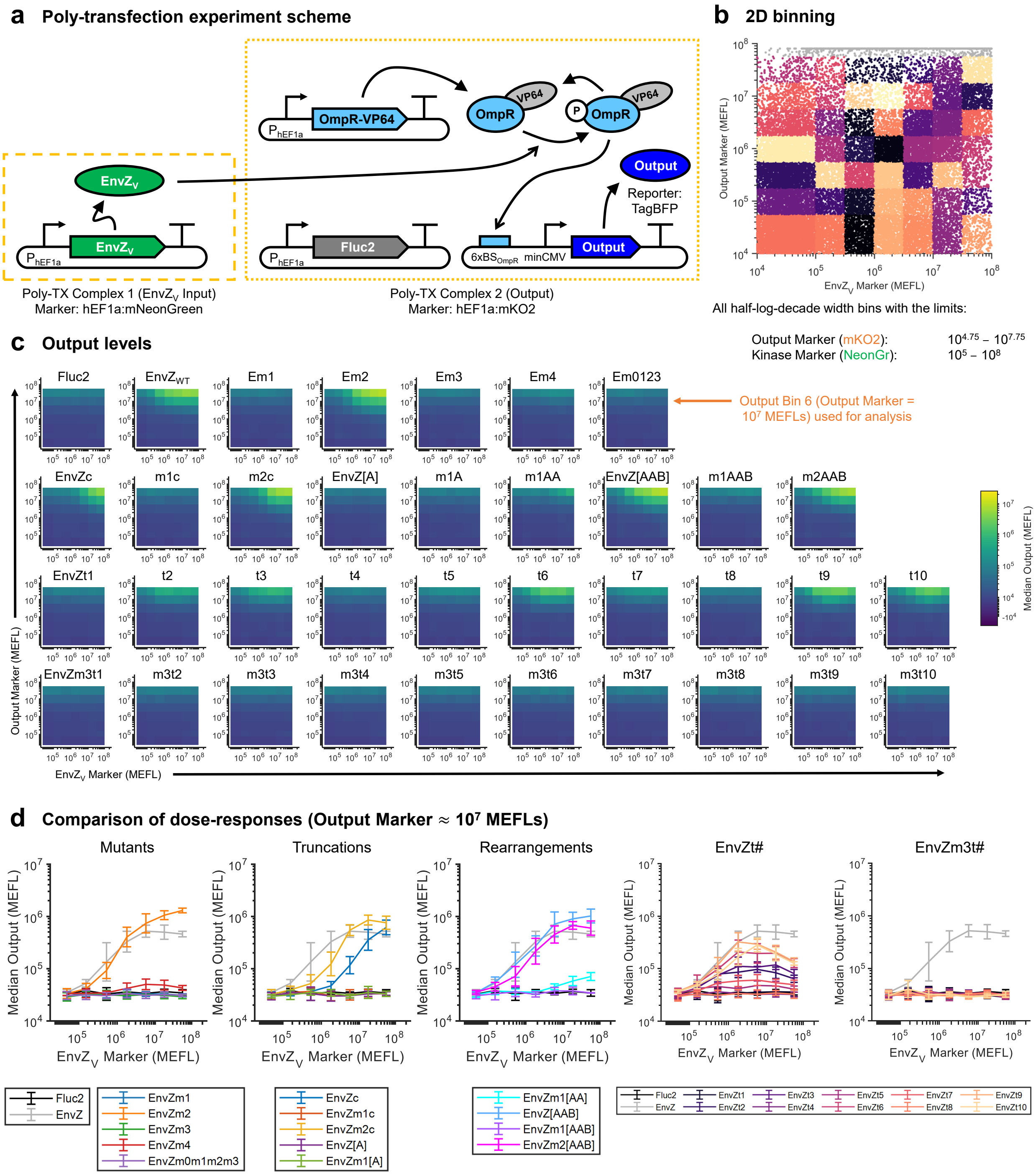


**Supplementary Figure 5: Test of EnvZ variants for OmpR-VP64 phosphorylation and output activation**.

**Supplementary Figure 5:** (Previous page.) **(a)** Schematic for poly-transfection experiment. EnvZ variants ($EnvZ_V$) were delivered in separate complexes to measure the dose response of the OmpR-driven output to $EnvZ_V$. OmpR-VP64 binds to DNA with ~20-fold higher affinity when phosphorylated (P-OmpR-VP64) than when not[50] (see also Supplementary Figure 17). **(b)** 2D binning scheme for analysis of the poly-transfection data. The decoupled delivery of the $EnvZ_V$ (as indicated by its marker) and the other plasmids (as indicated by the output marker) yield cells with various combinations of DNA dosages of each. **(c)** Median output expression level per bin for each variant tested. Output Bin 6 (Output Marker = $\sim 10^7$ MEFLs) was used to draw $EnvZ_V$-to-output dose-response curves in Panel (d) & Figure 2. **(d)** Dose responses of output to each EnvZ variant. Wild-type (WT) EnvZ and the negative control (the luminescent protein Fluc2) are included in each graph for comparison. All data was collected 48 hours post-transfection in HEK-293FT cells. Heatmaps represent the mean of measurements from three experimental repeats. Errorbars represent the mean ± s.d. of measurements from three experimental repeats. All data was collected 48 hours post-transfection in HEK-293FT cells.

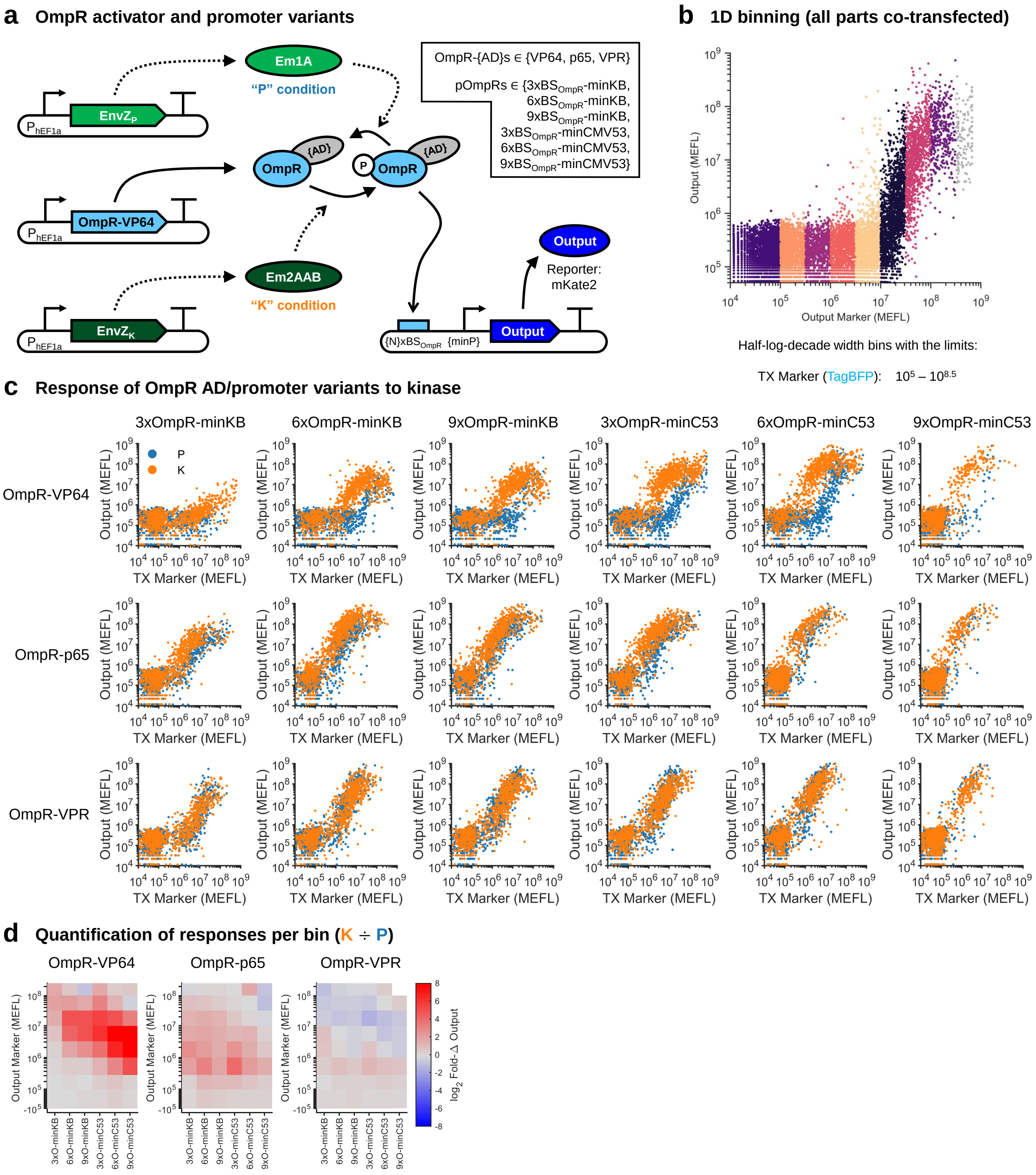


**Supplementary Figure 6: Characterization of OmpR activators and OmpR-activated promoters.**

**Supplementary Figure 6:** (Previous page.) **(a)** Schematic of experiment design; Either EnvZm1[A] or EnvZm2[AAB] was added, the former being a very weak/inactive phosphatase ("P" condition) and the latter a kinase ("K" condition) for OmpR. OmpR was fused to VP64, NF-$\kappa$B p65, or VPR activation domains. OmpR promoters were constructed with 3, 6, or 9 binding sites for OmpR upstream of a minimal promoter. The minimal promoter was either minCMV or minKB (*i.e.* YB_TATA)[22]. **(b)** 1D binning on the transfection marker. **(c)** Transfection scatterplots with data overlaid from each promoter/activator variant when co-transfected with EnvZm1[A] ("P") or EnvZm2[AAB] ("K"). The poor responsiveness to kinase for the p65 and VPR fusions likely results from the resulting protein being too strong of an activator, even when not phosphorylated. The 9xOmpR-minCMV promoter appeared to cause some toxicity, resulting in a reduced number of transfected cells. **(d)** Fold-change from "P" to "K" condition per bin for each combination of activator and promoter. OmpR-VP64 showed higher fold-changes in combination with promoters bearing more OmpR binding sites. OmpR-p65 had only modest fold-changes when used with weaker promoters and at lower transfection marker bins. OmpR-VPR had essentially no fold-change and possibly a minor decrease in expression at higher transfection marker bins. Based on this data, we used OmpR-VP64 and 6xOmpR-minCMV as the activator-promoter combination in all other experiments. All data was collected 48 hours post-transfection in HEK-293FT cells.

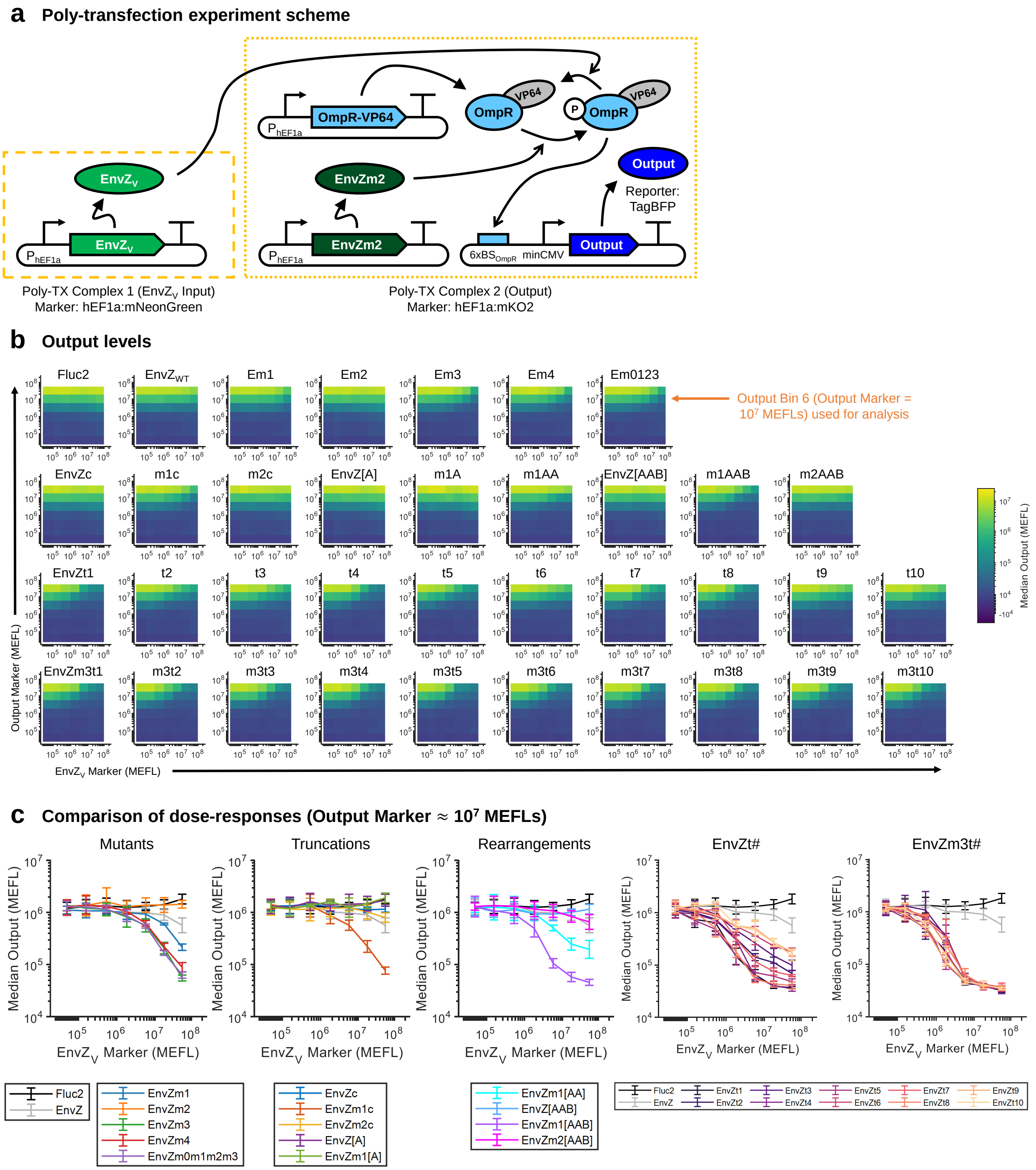


**Supplementary Figure 7: Test of EnvZ variants for OmpR-VP64 dephosphorylation and output deactivation.** **(a)** Schematic for poly-transfection experiment. Similar to Supplementary Figure 5, but EnvZm2 is expressed constitutively to generate P-OmpR-VP64 for dephosphorylation. **(b-c)** Similar analysis of 2D binning as in Supplementary Figure 5c-d. All data was collected 48 hours post-transfection in HEK-293FT cells.

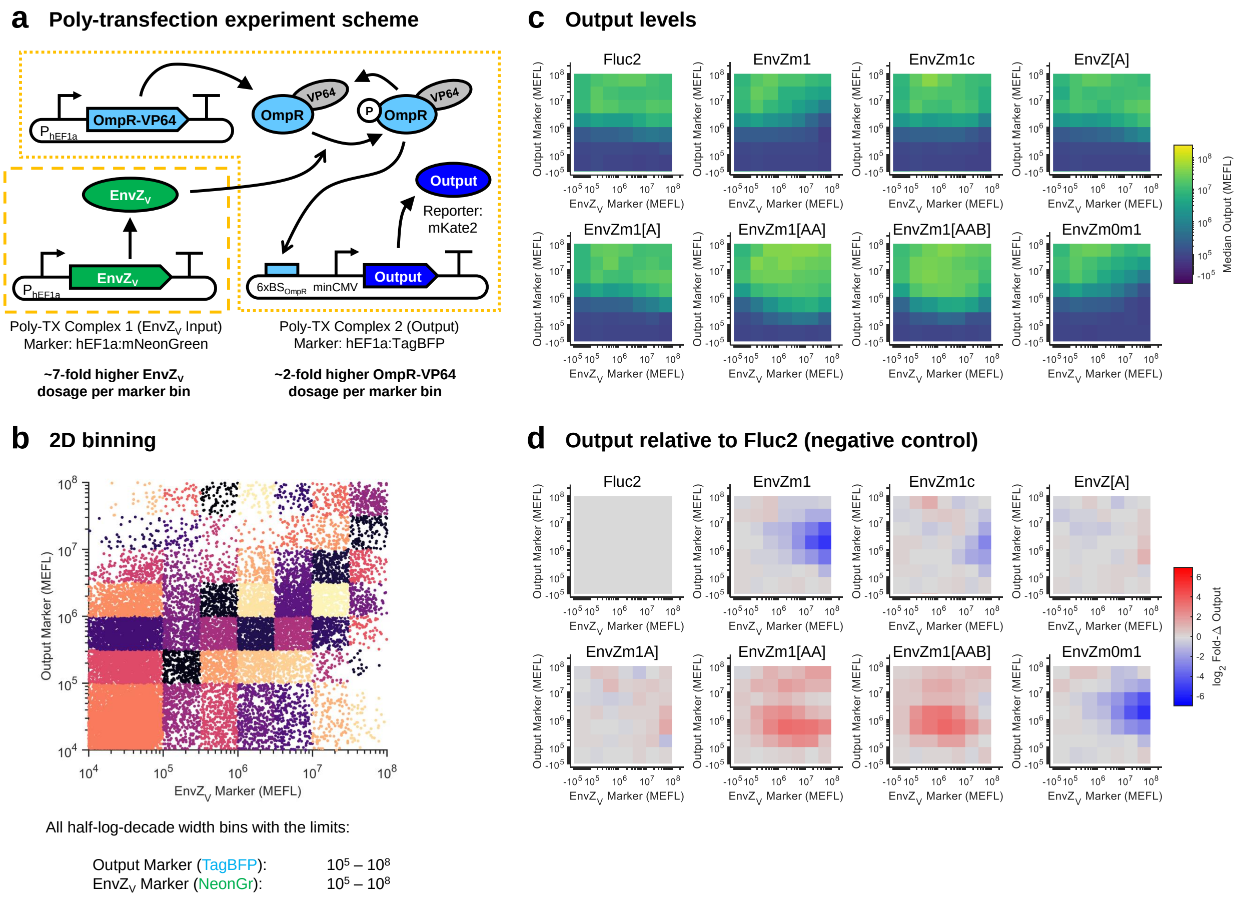


**Supplementary Figure 8: Evaluation of EnvZ variants at higher dosages**. **(a)** Schematic for poly-transfection experiment. Compared to the experiment in Figure 2, the ratio of EnvZ$_V$ to Marker in Complex 1 was ~7-fold higher, and the relative amount of OmpR-VP64 in Complex 2 was ~2-fold higher. **(b)** 2D binning scheme to analyze poly-transfection data. **(c)** Median output expression level per bin for each variant tested. **(d)** Fold-change in median expression level for each bin compared to the equivalent system expressing Fluc2 in place of EnvZ$_V$. All data was collected 48 hours post-transfection in HEK-293FT cells.

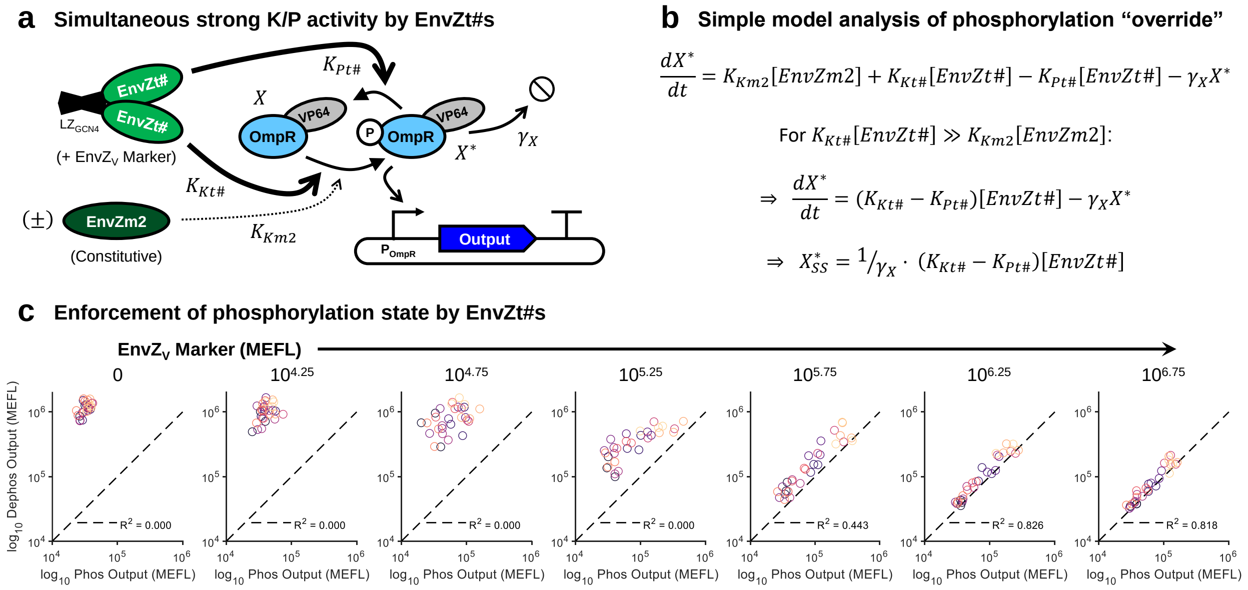


$$\frac{dX^*}{dt} = K_{Km2}[EnvZm2] + K_{Kt\#}[EnvZt\#] - K_{Pt\#}[EnvZt\#] - \gamma_X X^*$$

For $K_{Kt\#}[EnvZt\#] \gg K_{Km2}[EnvZm2]$:

$$\Rightarrow \frac{dX^*}{dt} = (K_{Kt\#} - K_{Pt\#})[EnvZt\#] - \gamma_X X^*$$

$$\Rightarrow X^*_{SS} = {}^1/_{\gamma_X} \cdot (K_{Kt\#} - K_{Pt\#})[EnvZt\#]$$

**Supplementary Figure 9: Enforcement of OmpR-VP64 phosphorylation level by bifunctional EnvZ variants**. **(a)** Schematic illustrating how simultaneous strong phosphorylation and dephosphorylation of OmpR-VP64 overrides phosphorylation from other sources (here the kinase EnvZm2). At a high level, the regulation of RR phosphorylation by a bifunctional HK is an example of feedforward control. **(b)** Simplified mathematical analysis of the system in Panel (a). If the reaction rates of OmpR phosphorylation and dephosophorylation are high enough, then rates corresponding to other kinases/phosphatases become effectively nullified and the output level is set by the bifunctional protein. In real systems, bifunctionality of the HK has been shown to impart insulation to cross-talk by other TCS pathways[19,20,106,107]. **(c)** Comparison of median output levels in response to activation and deactivation by EnvZt# variants in the absence and presence of EnvZm2, respectively. The plots show data from bins with increasing concentrations of each EvnZt# (from left to right, 0 to max EnvZt#), corresponding to the left two plots shown in Figure 2c. $R^2$ was computed for the 1:1 dashed line. Each circle represents a measurement from one experimental repeat. All data was collected 48 hours post-transfection in HEK-293FT cells.

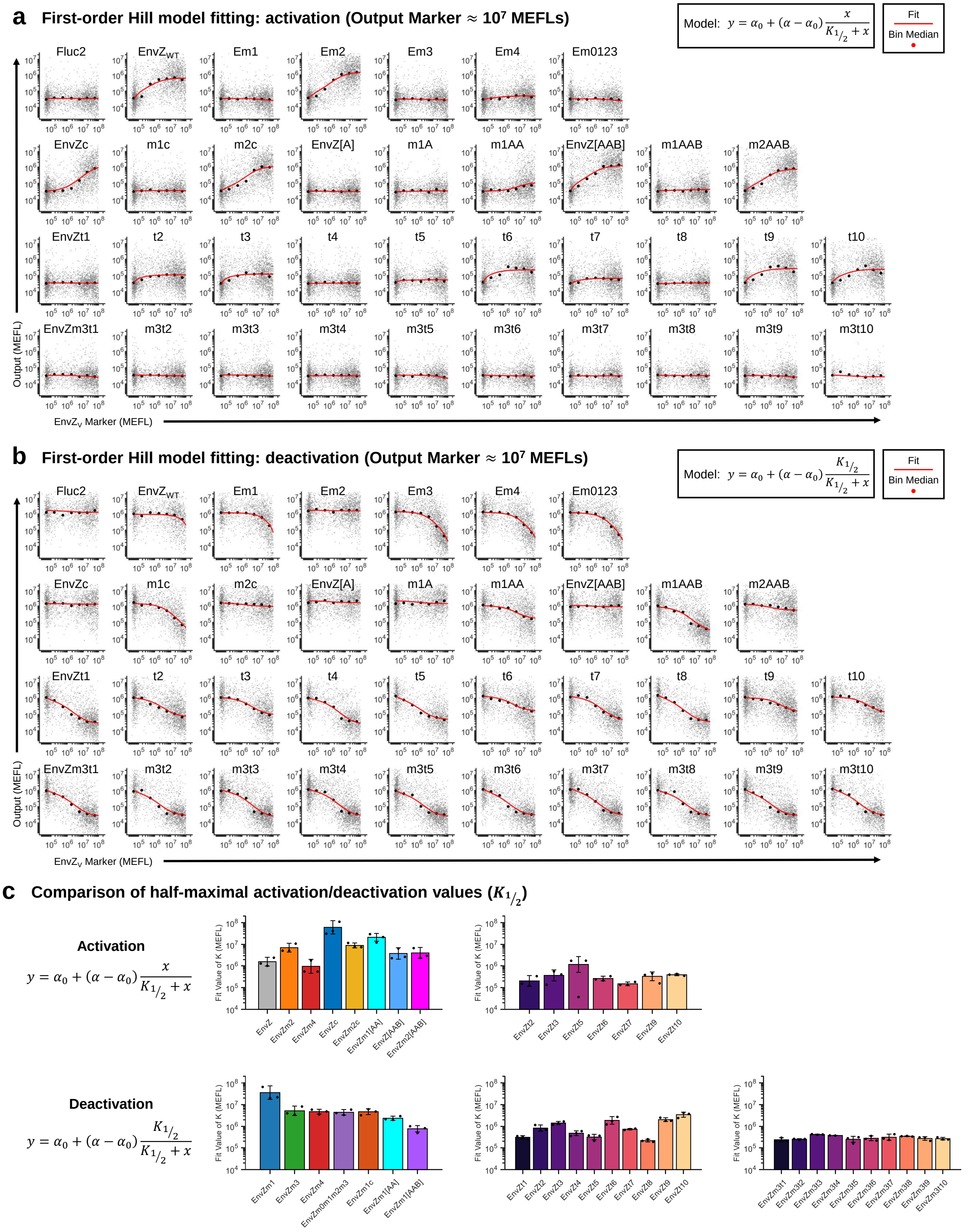


**Supplementary Figure 10: Fitting half-maximal response coefficients for activation and deactivation by each EnvZ variant**

**Supplementary Figure 10:** (Previous page.) **(a)** Fitting of activation functions. The model is shown in the top right and is a simple Hill model of activation corresponding to Equation (7) in Methods. The data plotted are extracted from the bins plotted in Supplementary Figure 5d, with individual cells shown as points and the bin medians and fit overlaid. Data is representative from the first of three experimental repeats. **(b)** Similar to Panel (a) but for the deactivation functions corresponding to Equation (8) in Methods and the data plotted in Supplementary Figure 7c. **(c)** Comparison of $K_{1/2}$ values fit for variants that showed activation (top) or deactivation (bottom). A lower value of $K_{1/2}$ indicates more potent activity by an enzyme. Errorbars indicate the mean ± s.d. of fit values for three experimental repeats.

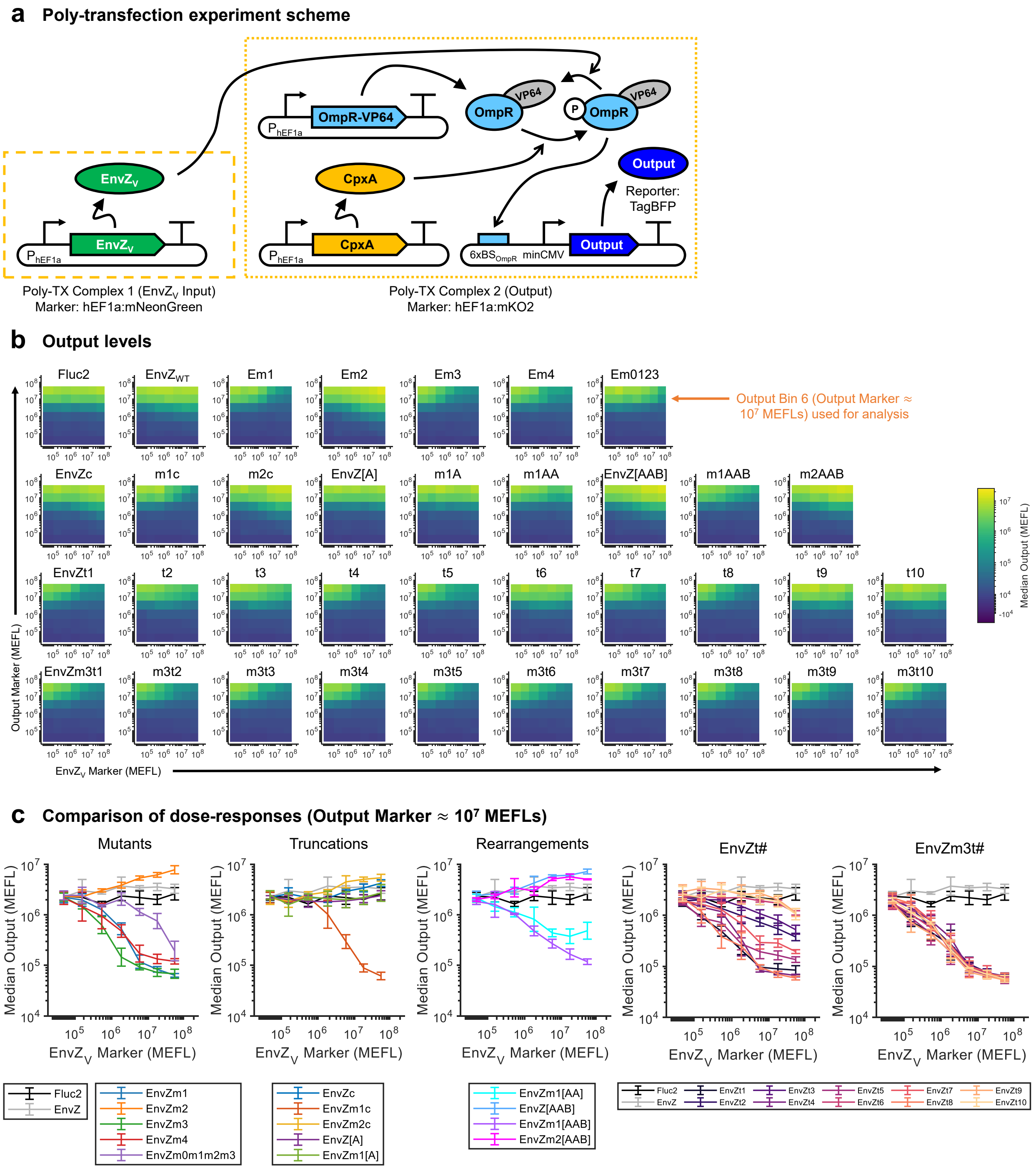


**Supplementary Figure 11: Dephosphorylation of CpxA-phosphorylated OmpR-VP64 by EnvZ variants**. **(a-c)** Analogous experiment to Figure 7, but with CpxA expressed instead of EnvZm2 to generate P-OmpR. CpxA has mild off-target phosphorylation of OmpR[20], and so can be used to avoid heterodimerization between EnvZ variants and EnvZm2. All data was collected 48 hours post-transfection in HEK-293FT cells.

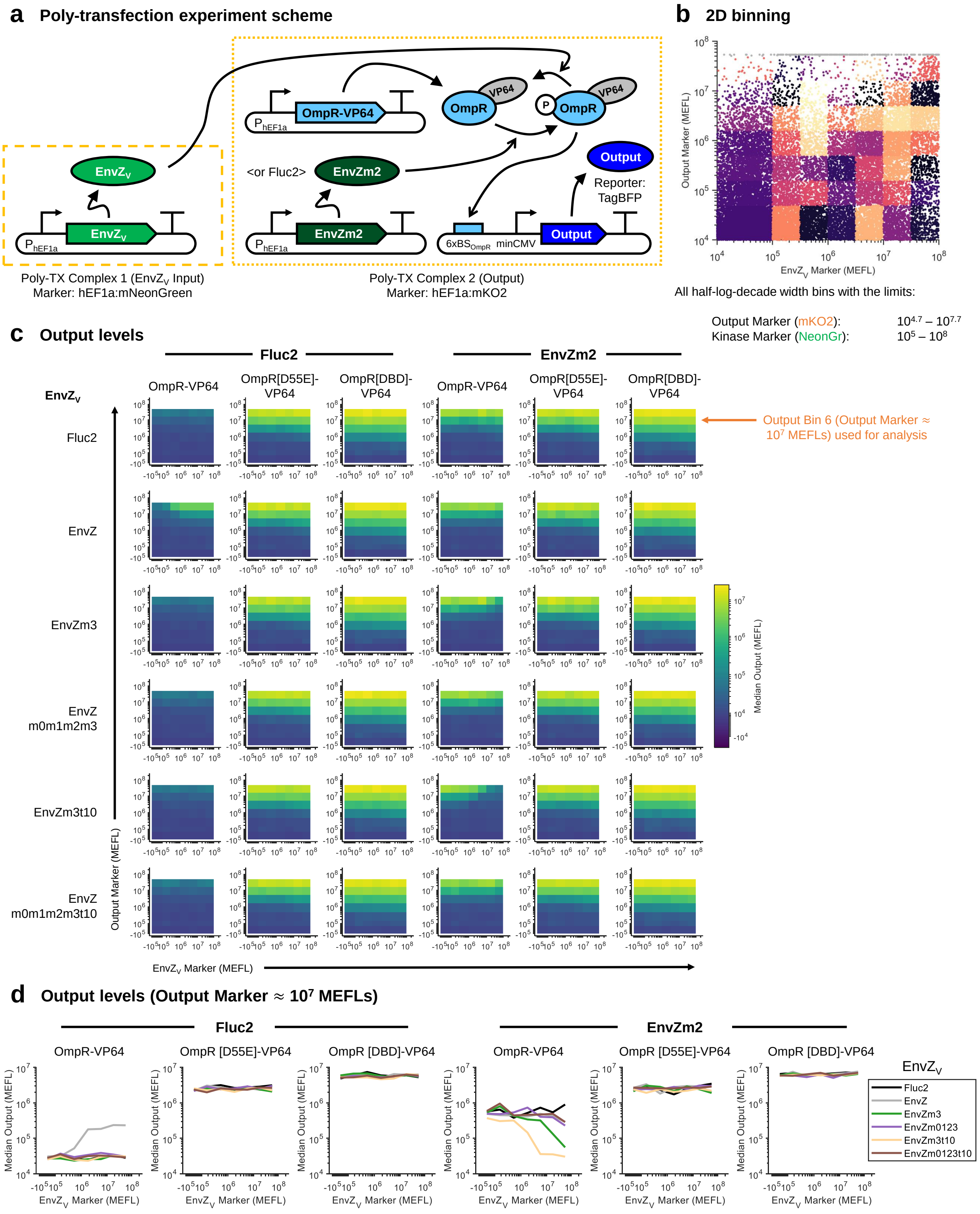


**Supplementary Figure 12: Additional controls to validate *in vivo* phosphatase activity.**

**Supplementary Figure 12:** (Previous page.) **(a)** Schematic of poly-transfection experiment. EnvZ variants (including Fluc2 as a negative control for phosphorylation) were delivered in a separate complex as OmpR-VP64, an OmpR-driven reporter plasmid, and either Fluc2 or the kinase EnvZm2. Two constitutive forms of OmpR (OmpR[D55E] and OmpR[DBD]) were used as controls for EnvZ-binding competent but phosphorylation-independent activation and EnvZ-binding incompetent activation, respectively. **(b)** 2D binning scheme for analysis of the poly-transfection data. **(c)** Median output expression level per bin for each variant tested. Output Bin 6 (Output Marker = $\sim 10^7$ MEFLs) was used to draw $\text{EnvZ}_V$-to-output dose-response curves in Panel (d). **(d)** Dose-responses of OmpR-driven output to EnvZ variants in the presence/absence of EnvZm2 kinase. With or without the kinase, constitutive forms of OmpR-VP64 strongly activate output expression regardless of the dosage of any given EnvZ variant, indicating that sequestration of OmpR does not causes output reduction at the concentrations tested (which match those in most experiments in the study except Supplementary figure 8, in which the dosages were purposefully elevated further). Only wild-type EnvZ activates wild-type OmpR-VP64, and knockout of phosphatase activity in EnvZm3t10 (our selected phosphatase variant for the majority of experiments with a phosphatase) specifically prevents deactivation of OmpR-driven expression. Thus, the deactivation observed can be very likely attributed to dephosphorylation of OmpR-VP64, rather than sequestration of OmpR or inhibition of binding between OmpR and the kinase.

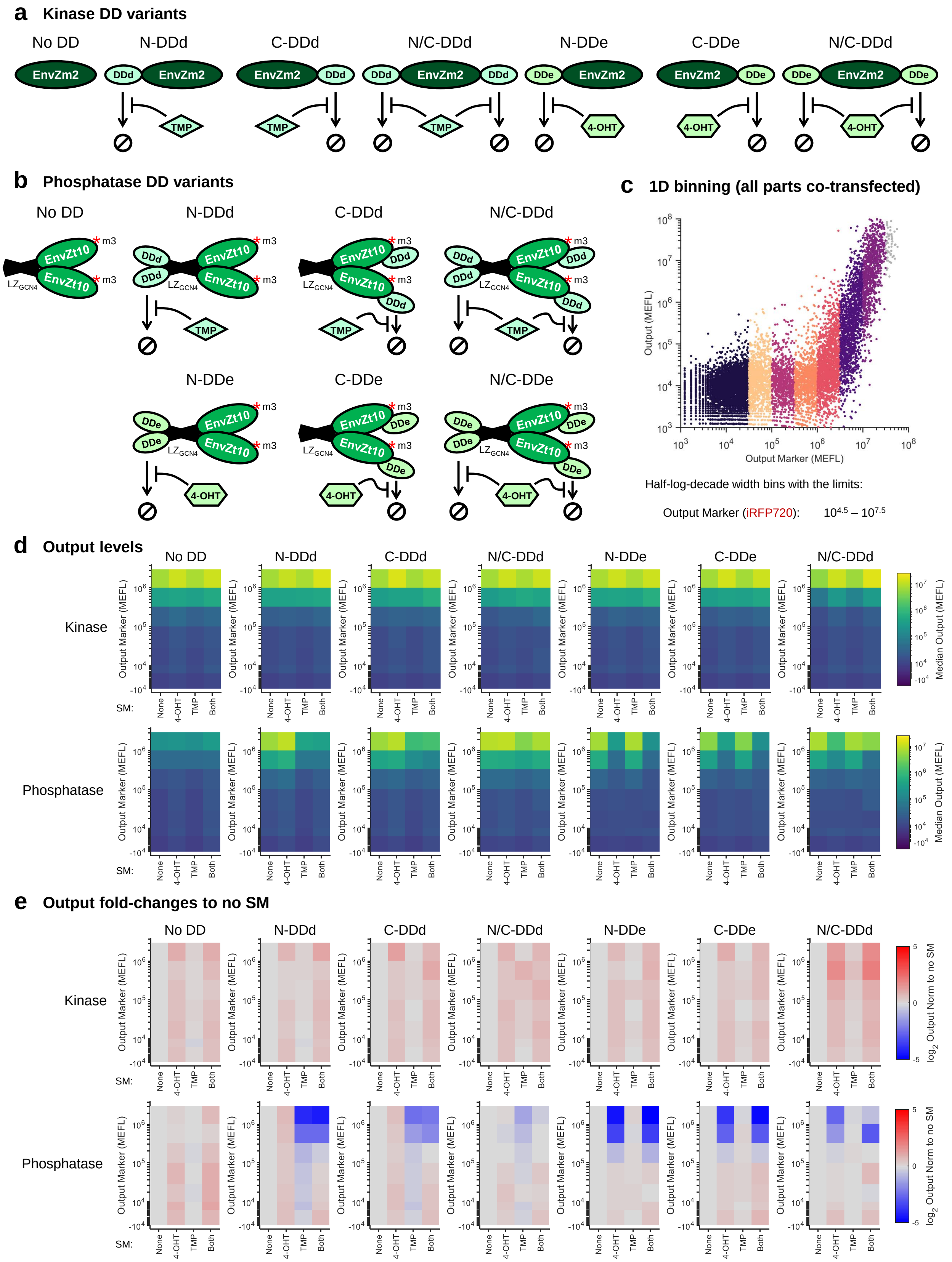


**Supplementary Figure 13: Fusions of inducible degradation domains to EnvZ kinase/phosphatase**

**Supplementary Figure 13:** (Previous page.) **(a)** Variants of DDd[45] and DDe[46] fused to the N-, C-, or both termini of EnvZm2 (kinase). DDd and DDe are unstable until bound by their respective inducer molecules, trimethoprim (TMP) and 4-hydroxytamoxifen (4-OHT). **(b)** Variants of DDd and DDe fused to EnvZm3t10 (phosphatase). **(c)** 1D binning of a co-transfection of hEF1a:OmpR-VP64, one of the DD-EnvZm2 variants or one of the DD-EnvZm3t10 variants + EnvZm2, an OmpR-driven fluorescent reporter, and the constitutive transfection marker used for binning. **(d)** Median output level per bin for all combinations of neither, one, or both inducer molecules. The top row is for the DD-kinases and the bottom for the DD-phosphatases. **(e)** Fold-changes per bin for each DD-kinase and DD-phosphatase variant to the sample with no small molecule(s) added. All data was collected 48 hours post-transfection in HEK-293FT cells.

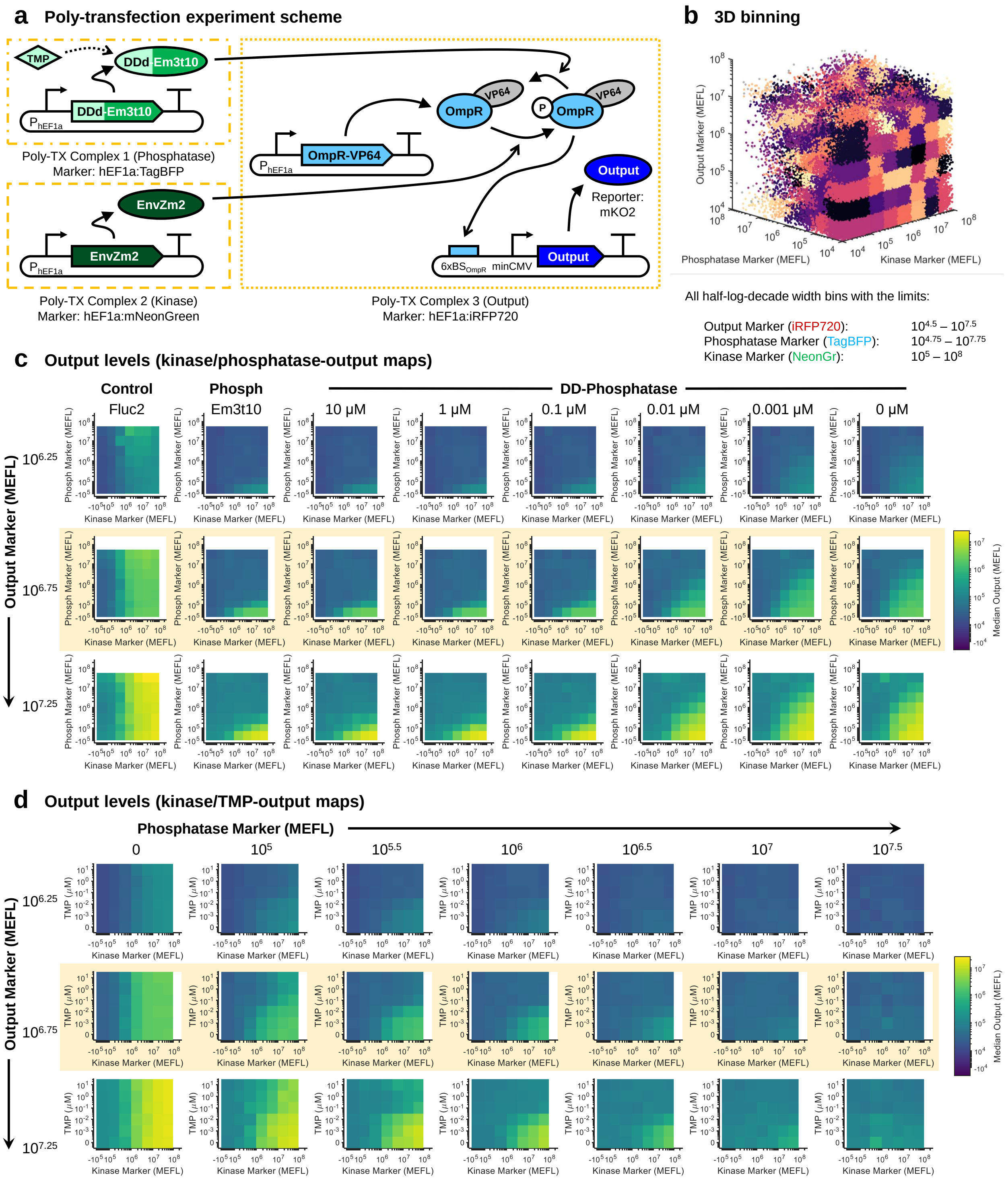


**Supplementary Figure 14: Tuning expression with kinase and TMP-inducible phosphatase**

**Supplementary Figure 14:** (Previous page.) **(a)** Schematic for poly-transfection experiment. TMP stabilizes DDd-EnvZm3t10. **(b)** 3D binning scheme to analyze poly-transfection data. **(c)** Median output expression across all Kinase/Phosphatase Marker bins and at select Output Marker bins. The highlighted row corresponds to the data in Supplementary Figure 15. 'Control' refers to a sample with Fluc2 added in place of the phosphatase. 'Phosph' refers to a sample expressing EnvZm3t10 (phosphatase) not fused to a DD. 'DD-phosphatase' refers to samples with DDd-EnvZm3t10 and the indicated concentration of TmP in the media. **(d)** A re-arrangement of the DDd-EnvZm3t10 samples in Panel (c) to highlight the 2D response to kinase/TMP inputs per Phosphatase Marker bin. Each heatmap in (c) and (d) shows the mean of measurements from three experimental repeats. All data was collected 48 hours post-transfection in HEK-293FT cells.

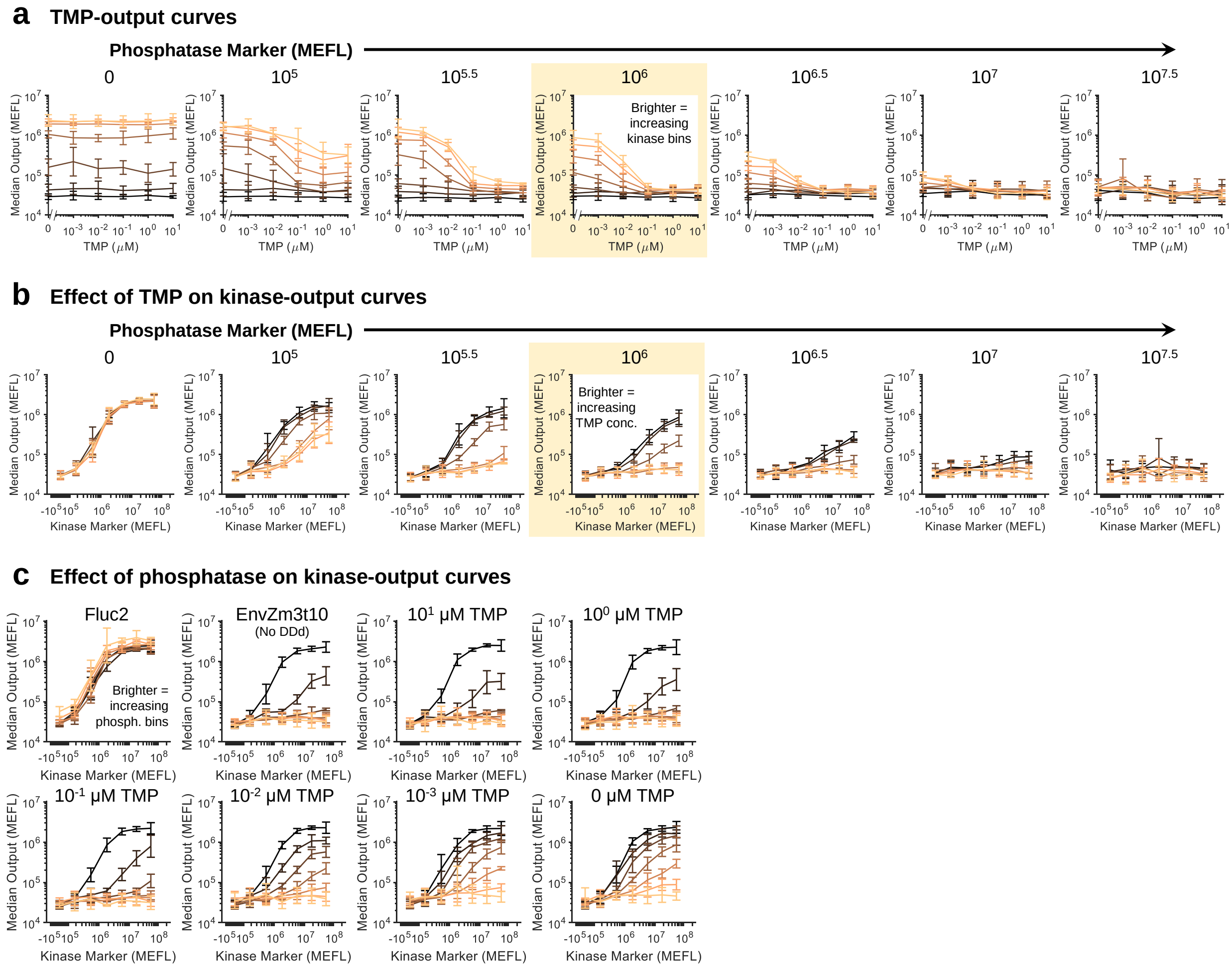


**Supplementary Figure 15: Comparison of dose-response curves for TMP-inducible phosphatase circuit**. **(a-b)** Dose-responses of output to TMP (a) or kinase (b) per Phosphatase Marker bin. Brighter lines indicate bins with more kinase (a) or TMP (b). The highlighted plots are shown in Figure 3c. **(c)** Dose responses to kinase per phosphatase variant. Brighter lines indicate bins with more phosphatase. Note that 'Fluc2' corresponds to a control with Fluc2 expressed in place of a phosphatase. In all plots, the errorbars represent the mean ± s.d. of measurements from three experimental repeats.

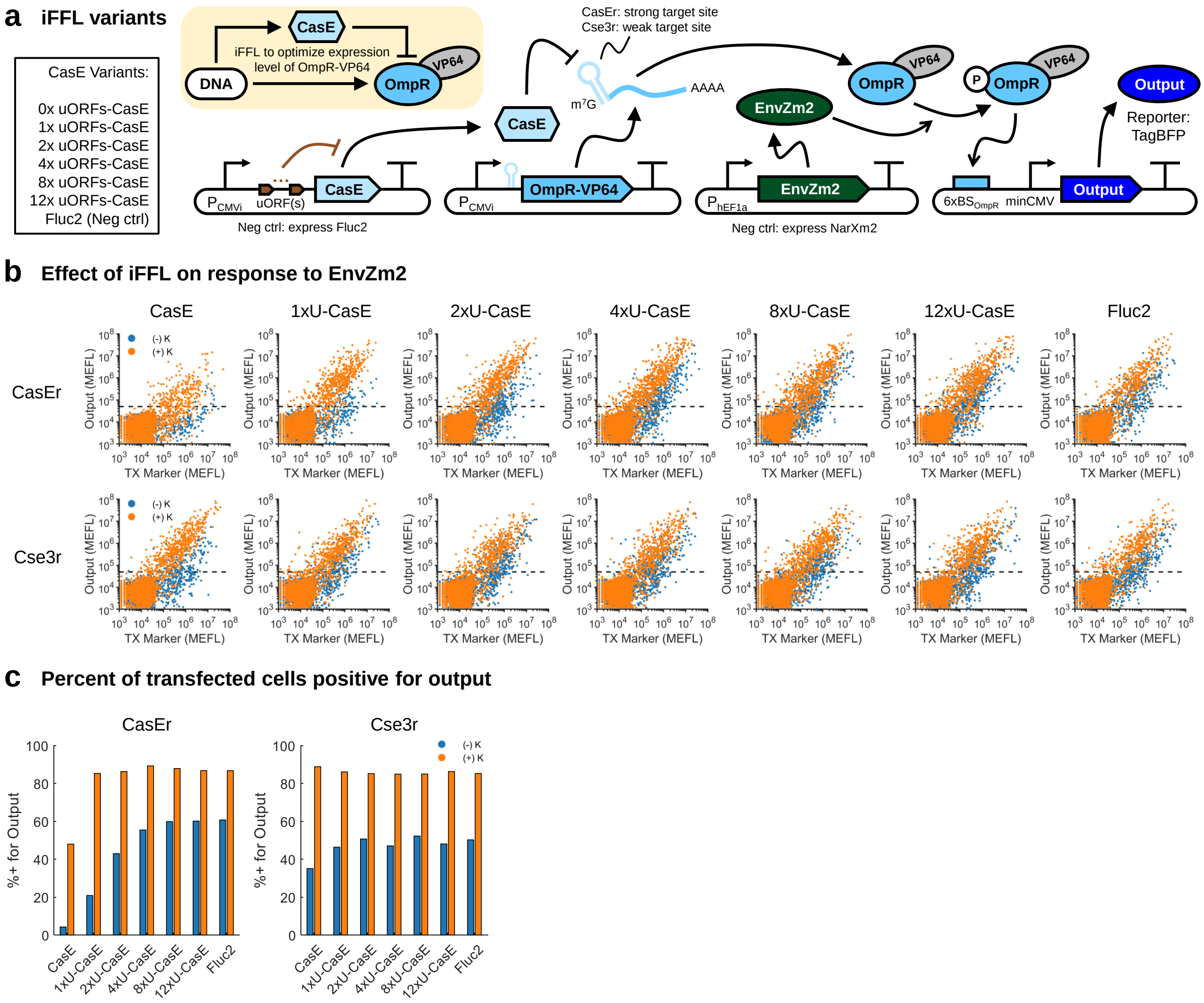


**Supplementary Figure 16: Feedforward control to optimize the level of OmpR-VP64**. **(a)** An endoRNase-based incoherent feedforward loop (iFFL)[51] to optimize OmpR-VP64 levels above the $K_D$ of P-OmpR but below that of OmpR[50] (Supplementary Figure 17). CasE is a CRISPR endoRNase[108,109] that binds and cleaves a ~20 bp target site placed in the 5'UTR of OmpR-VP64. More uORFs[110] in the 5'UTR of CasE reduce its translation rate and thus expression level. **(b)** Transfection scatterplots for each iFFL variant with (+) and without (–) EnvZm2. In the absence of EnvZm2, Fluc2 is expressed instead. The dashed line indicates the threshold above which cells are 'positive' for the output reporter. Target sites for both CasE (CasEr) and Cse3 (Cse3r) were compared, the latter of which is also cleaved by CasE, though less efficiently[109]. Cse3r samples were inlcuded in case the knockdown of CasEr was too strong, though that ultimately was not an issue. **(c)** Percent of transfected cells positive for the output in all samples. Cells were gated as transfected if they were positive for the output or the constitutive transfection marker. Though the maximum difference in percent positive is seen with 1x uORFs-CasE, the reduced percent positive for CasE without uORFs indicates improved selectivity for output expression in the absence of kinase input, which is preferable for ensuring a strong 'off' state. All data was collected 48 hours post-transfection in HEK-293FT cells.

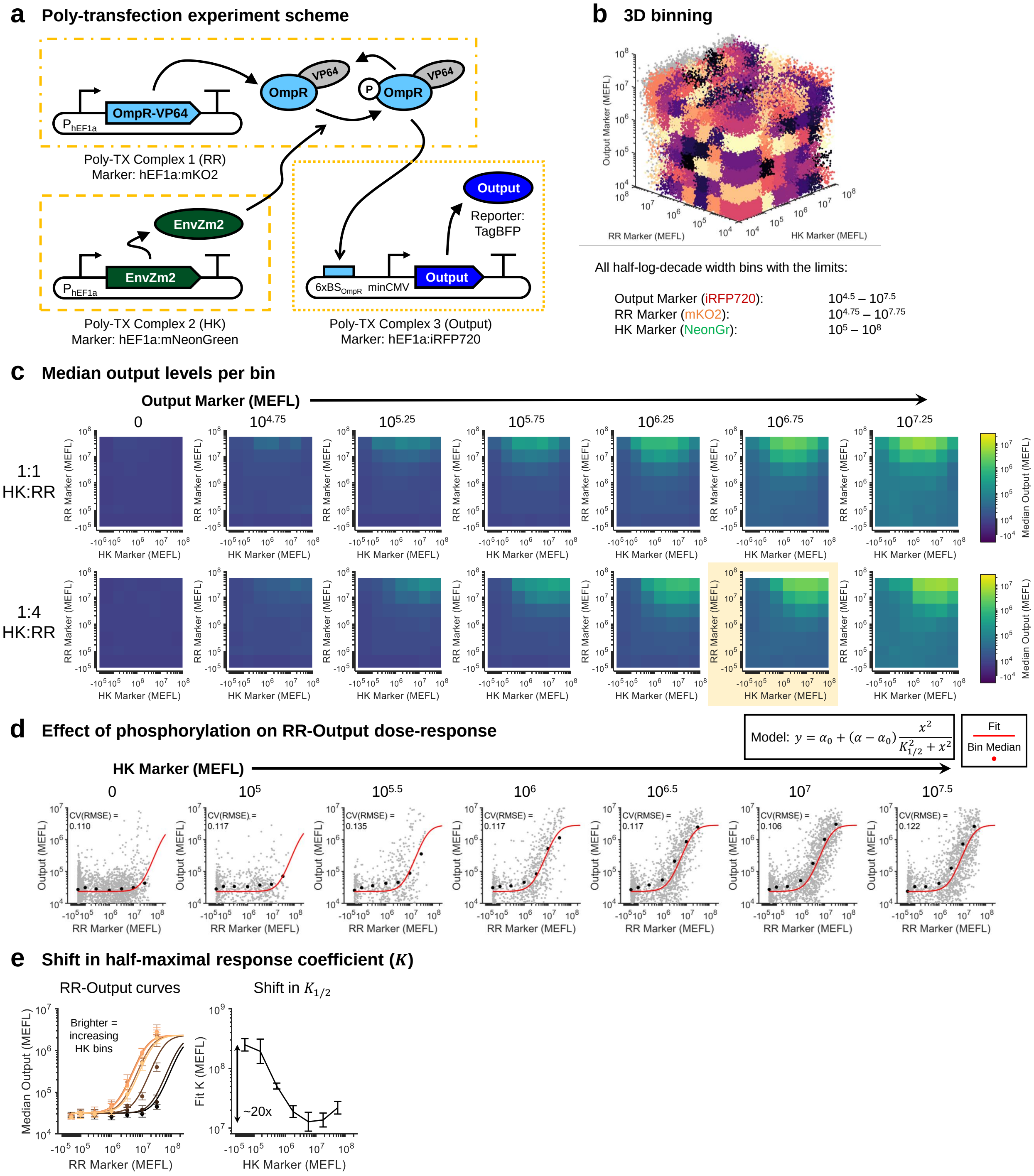


**Supplementary Figure 17: 2D EnvZ-OmpR dose-response curves.**

**Supplementary Figure 17:** (Previous page.) **(a)** Schematic for poly-transfection experiment. Here OmpR-VP64 was separated into a third DNA-lipid complex in order to separate its effect on output expression from that of increasing reporter plasmid. **(b)** 3D binning scheme to analyze poly-transfection data. **(c)** Median output level in all bins for two versions of the experiment: (top) equal plasmid dosages for EnvZ and OmpR-VP64 in their respective complexes and (bottom) 1:4 diluted EnvZ compared to OmpR-VP64. Relatively lower dosages of EnvZ avoid a knockdown of output expression at high EnvZ levels, likely due to sequestration of OmpR-VP64. Thus, the 1:4 ratio of HK:RR was used in other poly-transfection experiments. Data is the mean of measurements from three experimental repeats. **(d)** Fitting of OmpR-VP64 activation functions per level of EnvZ. The model is shown in the top right and is a simple Hill model of activation corresponding to Equation (6) in Methods. The cooperativity is assumed to be 2 because OmpR binds to DNA as a dimer[15,92]. The data plotted are extracted from the bins plotted in the highlighted heatmap in Panel (c), with individual cells shown as points and the bin medians and fit overlaid. All fit parameters except $K_{1/2}$ were forced to be the same across each level of EnvZ. Data is representative from the first experimental repeat. **(e)** Change in $K_{1/2}$ as a function of EnvZ dosage. All data was collected 48 hours post-transfection in HEK-293FT cells.

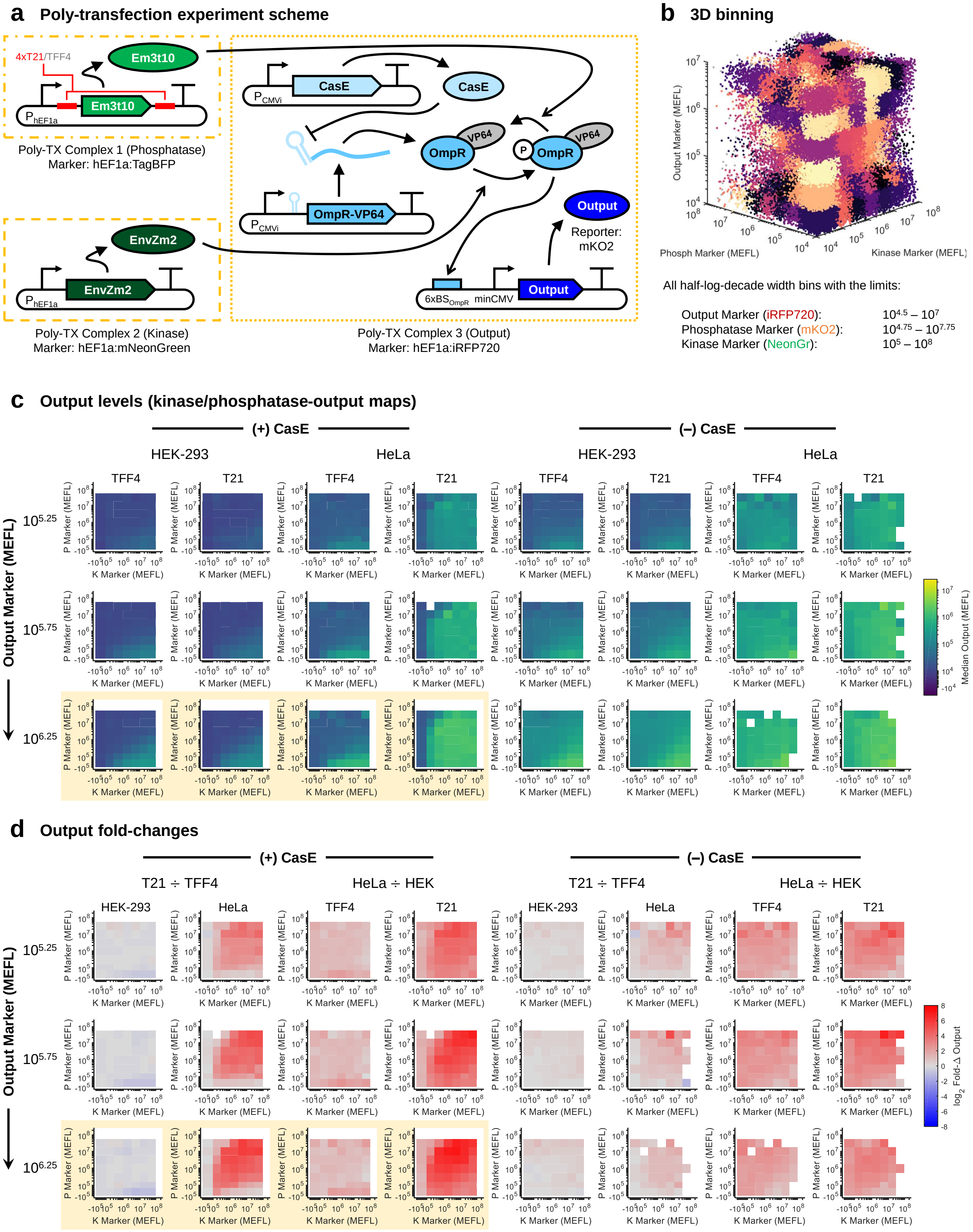


**Supplementary Figure 18: Extended data analysis of miR-21 sensor in HEK vs HeLa cells**

**Supplementary Figure 18:** (Previous page.) **(a)** Schematic for poly-transfection experiment. The CasE iFFL from Supplementary Figure 16 is incorporated into the output/OmpR-VP64 complex to optimize OmpR-VP64 levels. **(b)** 3D binning scheme to analyze poly-transfection data. **(c)** Median expression level per bin for all dosages of kinase and phosphatase inputs and select output marker bins. The expression levels in the absence of CasE are shown in the right grouping of plots. In the absence of CasE, Fluc2 was expressed instead. The highlighted plots correspond to line plots in Figure 4b – (+) and (–) phosphatase in that figure correspond to the phosphatase bins 1 and 6 (second-highest). The highest phosphatase bin was not used because the number of points was relatively sparse and thus noisy. **(d)** Fold-changes in output expression levels for the indicated comparisons. Fold-changes are improved when CasE is added compared to no CasE. All data was collected 48 hours post-transfection in HEK-293 or HeLa cells as indicated. Heatmaps show the mean of measurements from three experimental repeats.

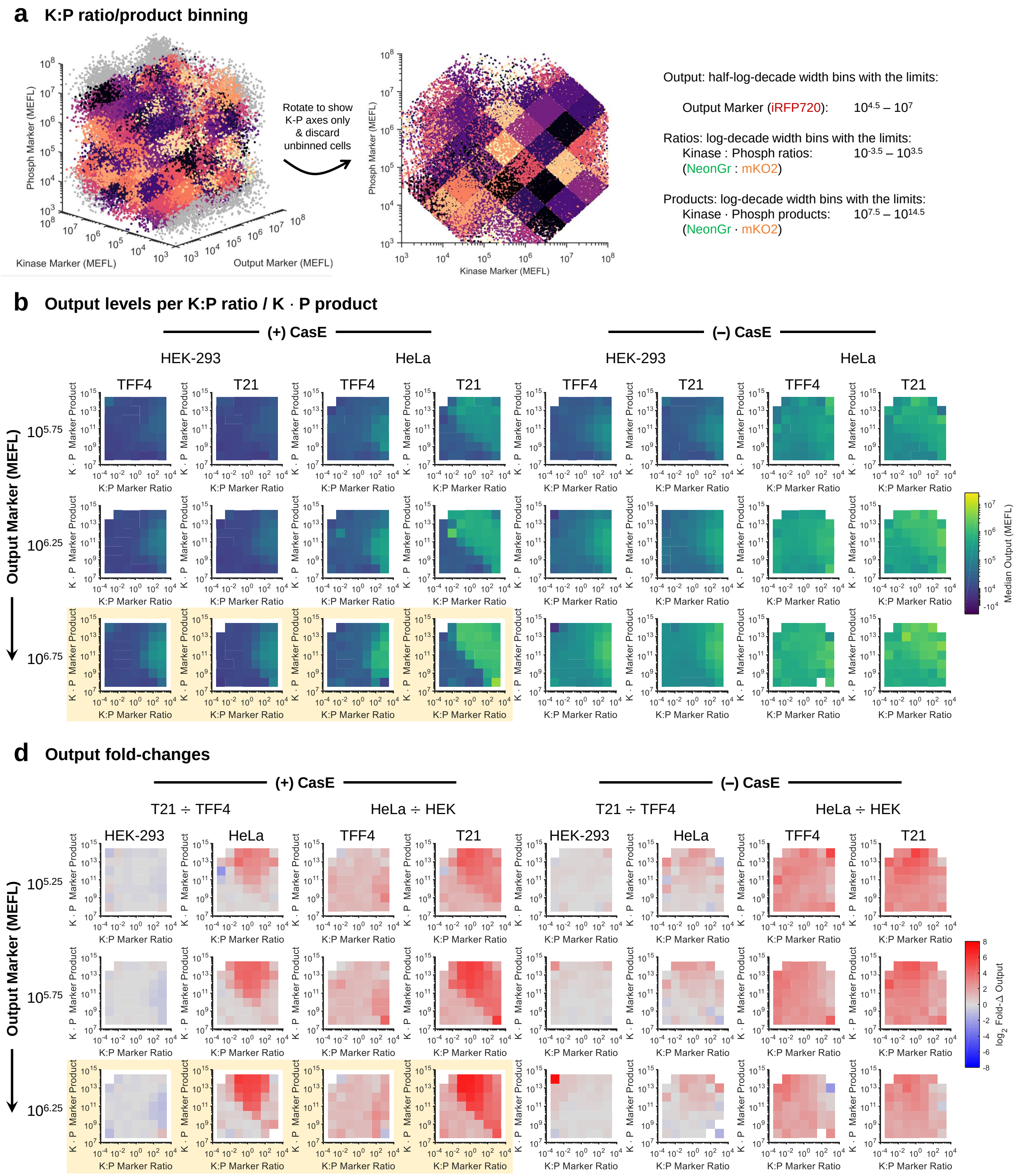


**Supplementary Figure 19: Ratiometric binning and analysis of miR-21 sensor.** **(a-d)** Similar analysis to Supplementary Figure 18, but with bins defined ratiometrically for the kinase and phosphatase markers, rather than in absolute terms. The product of kinase and phosphatase markers refers to bins that have the same kinase:phosphatase ratio but different total amounts of each.

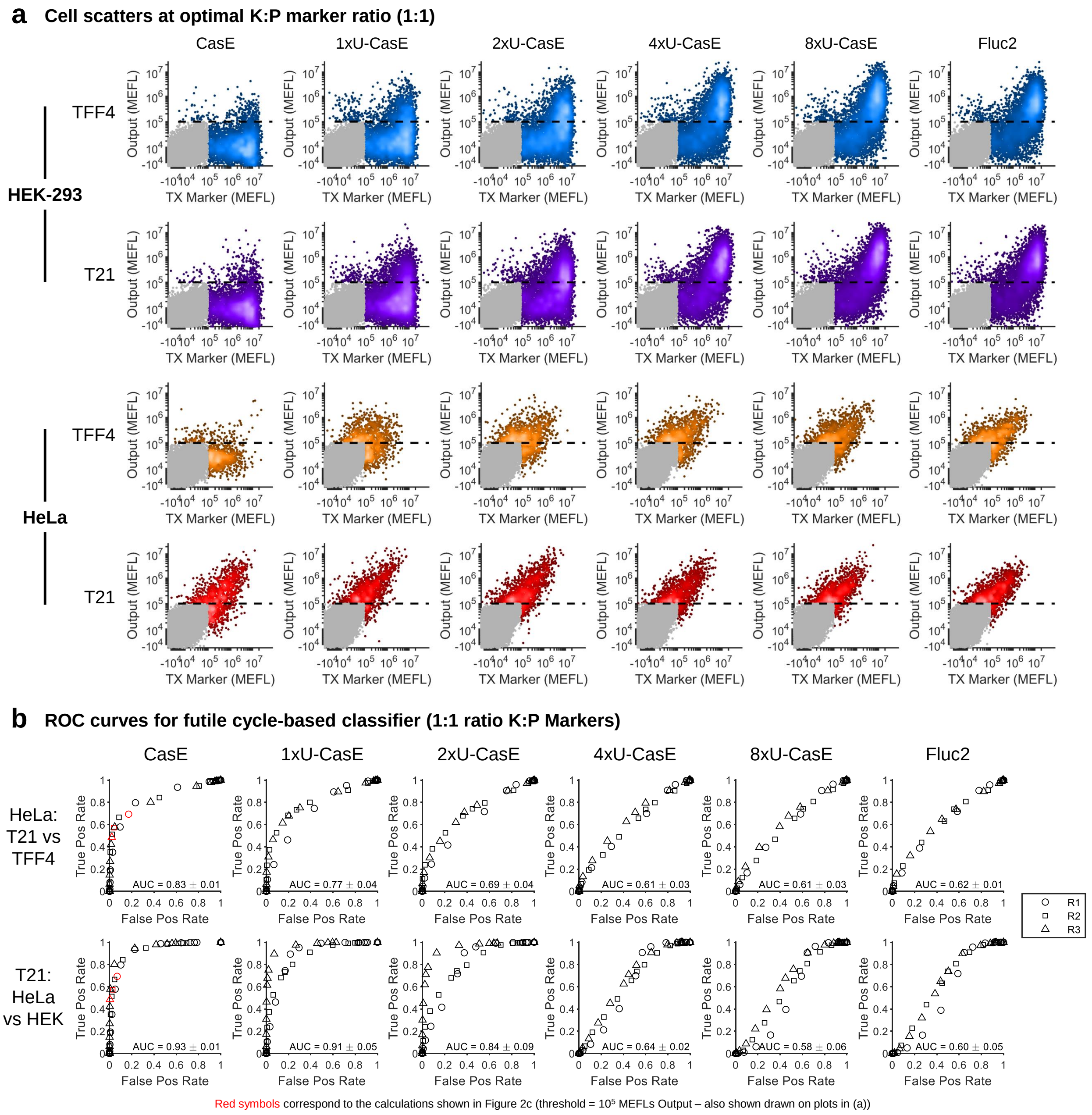


**Supplementary Figure 20: miR-21 sensing performance across classifier variants**. **(a)** Expression of output at optimal 'trajectories' subsampled from poly-transfection data (see Supplementary Figure 18). The ratio is 1:1:0.5 (Kinase Marker):(Phosphatase Marker):(Output Marker). Dashed lines indicate the threshold above which cells are considered positive for output (corresponding to the plots and percent positive graphs shown in Figure 4). Transfected cells are highlighted with the colors representing density. The data shown is representative from the first of three experimental repeats. **(b)** Receiver operating characteristic (ROC) curves for each sensor variant. True positive rate is the fraction of cells positive in the HeLa/T21 sample; false positive rate is the fraction of cells positive in the HeLa/TFF4 or HEK/T21 samples, as indicated. The red symbols correspond to the threshold shown in Panel (a): Output > $10^5$. The area under the curve (AUC) is given as the mean ± s.d. of measurements from three experimental repeats.

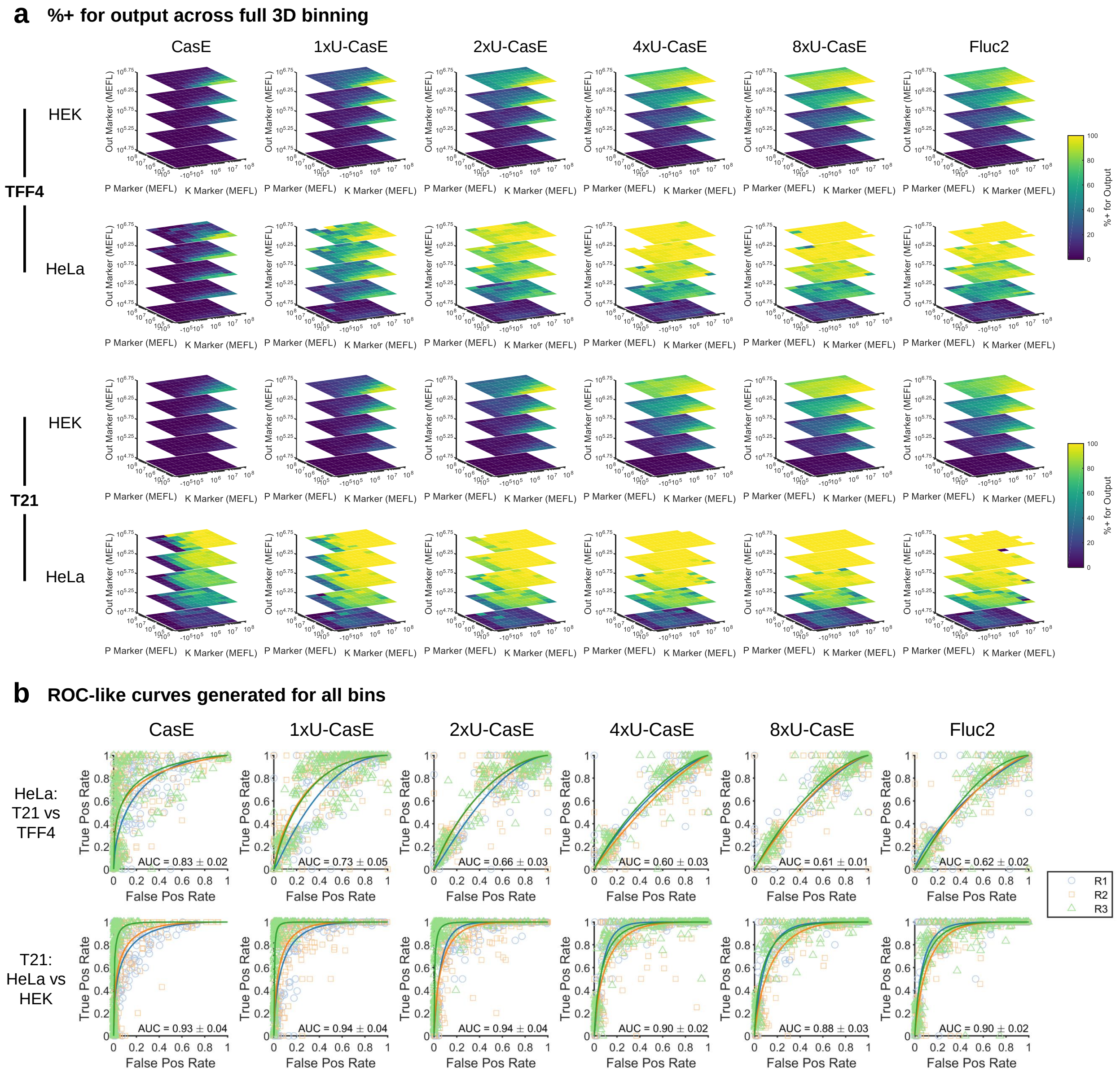


**Supplementary Figure 21: Comparison of miR-21 sensor performance across complete binned data**. **(a)** Percent of cells positive for output in each bin of the poly-transfection experiment (see Supplementary Figure 18). Values shown are the mean of three experimental repeats. **(b)** ROC-like curves generated by combining data from all bins for a given sensor variant onto one plot. ROC curves were generated by fitting the points with a bi-normal model. The AUC is given as the mean ± s.d. from three experimental repeats.

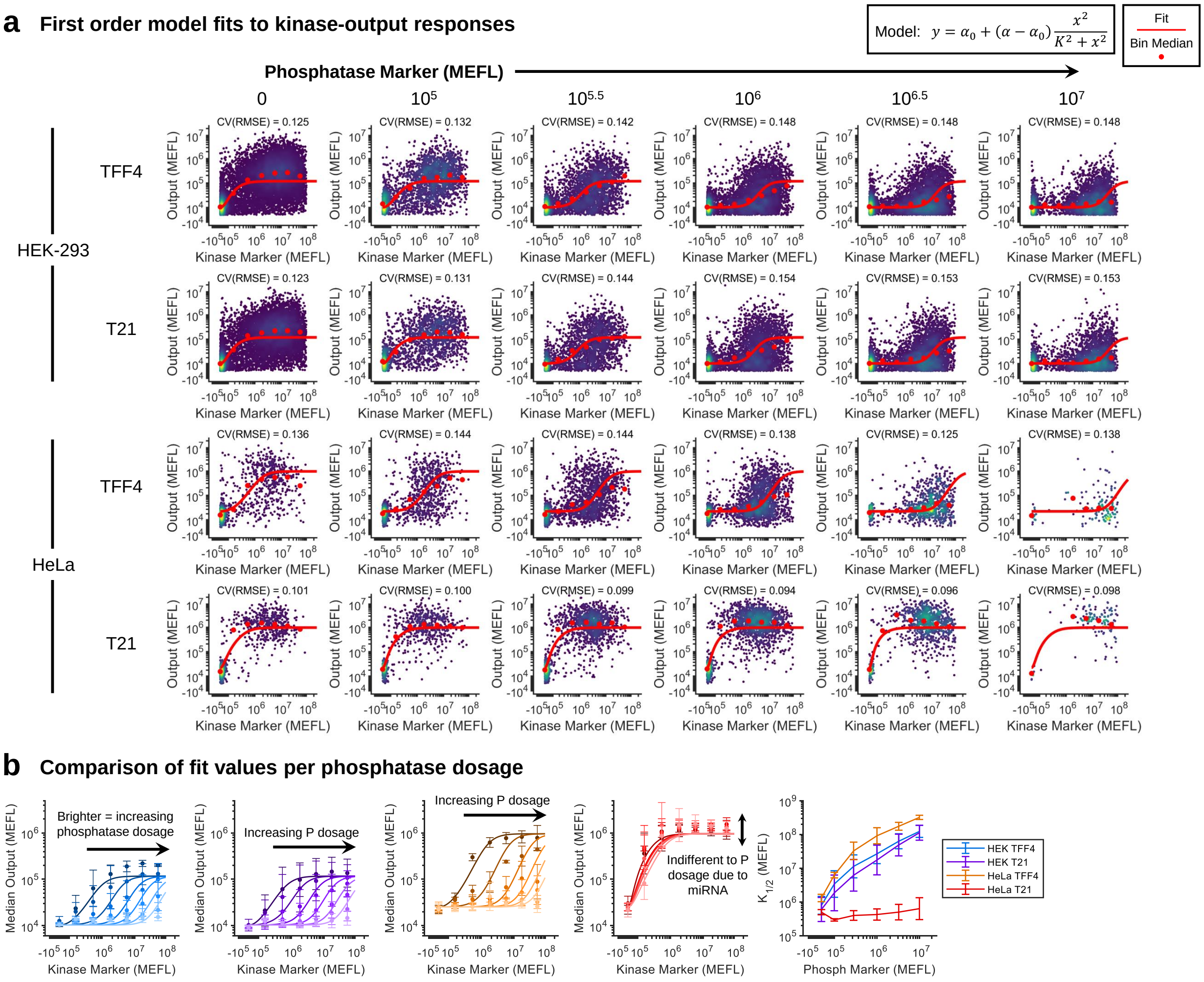


**Supplementary Figure 22: Desensitization of kinase-output curve to phosphatase levels in HeLa cells due to miR-21.** **(a)** Fitting of kinase (EnvZm2) activation functions per level of phosphatase (EnvZm3t10). The model is shown in the top right and is a simple Hill model of activation corresponding to Equation (7) in Methods. The cooperativity is assumed to be 2 because of OmpR dimerization once phosphorylated by EnvZm2 and the amount of P-OmpR-VP64 is assumed to be proportional to the level of kinase. The data plotted are extracted from the bins plotted in the highlighted heatmap in Panel (c), with individual cells shown as points and the bin medians and fit overlaid. All fit parameters except $K_{1/2}$ were forced to be the same across each level of phosphatase. Data is representative from the first experimental repeat. **(b)** Direct comparison of medians and fits for each phosphatase dosage, with the far-right plot showing the fit values of $K_{1/2}$ as a function of phosphatase dosage. Errorbars represent the mean ± s.d. of measurements from three experimental repeats.

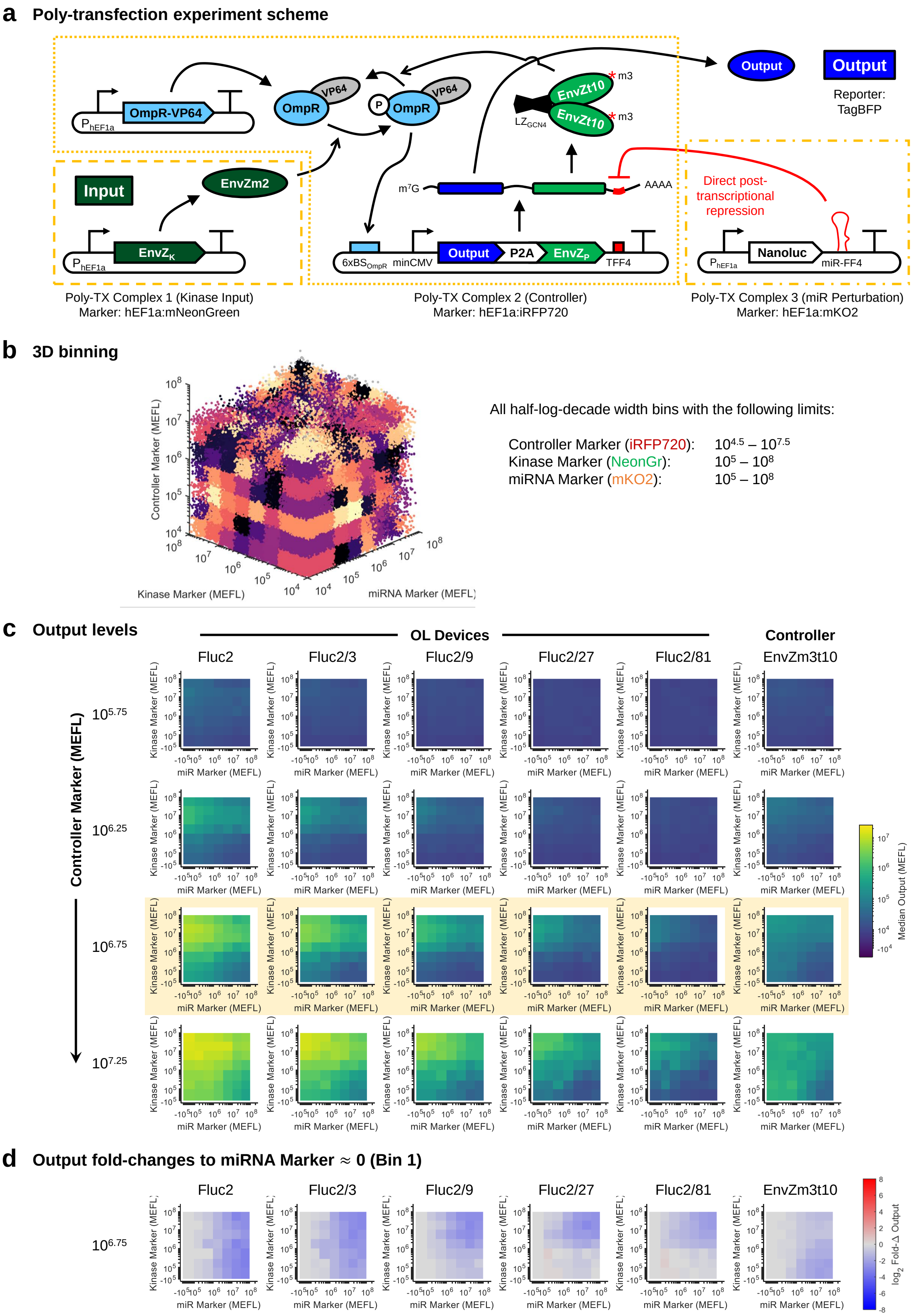


**Supplementary Figure 23: Extended analysis of OL variants with miR-FF4 perturbation**

**Supplementary Figure 23:** (Previous page.) **(a)** Schematic for poly-transfection experiment. miR-FF4 knocks down output expression by binding and cleaving a single target site in its 3'UTR. The output and phosphatase (EnvZm3t10) are 2A-linked to ensure equivalent transcriptional kinetics and co-knockdown by the miR. Separation of the kinase (EnvZm2) and miR-FF4 into separate lipid-DNA complexes enables simultaneous and independent measurement of their dose-responses on the output. **(b)** 3D binning scheme to analyze poly-transfection data. **(c)** Median levels across all Kinase Marker and miR Marker bins and select Controller Marker bins. OL refers to open loop versions of the controller, in which Fluc2 is expressed in place of the phosphatase. OL variants with reduced expression were created by diluting the output plasmid relative to other plasmids in Complex 2 by the indicate dilution factors (1:3, 1:9, 1:27, 1:81). The highlighted plots correspond to the data shown in Panel (d), Figures 5 & 6, and Supplementary Figures 27 & 29. These bins were chosen because they have enough Complex 2 to enable easily-detectable output reporter while not too much to cause substantial activation by unphosphorylated OmpR-VP64. **(d)** Fold-changes relative to miR Bin 1 (0 MEFLs) for the highlighted samples in Panel (c). All data was collected 48 hours post-transfection in HEK-293FT cells. Each heatmap in (c) and (d) shows the mean of measurements from three experimental repeats.

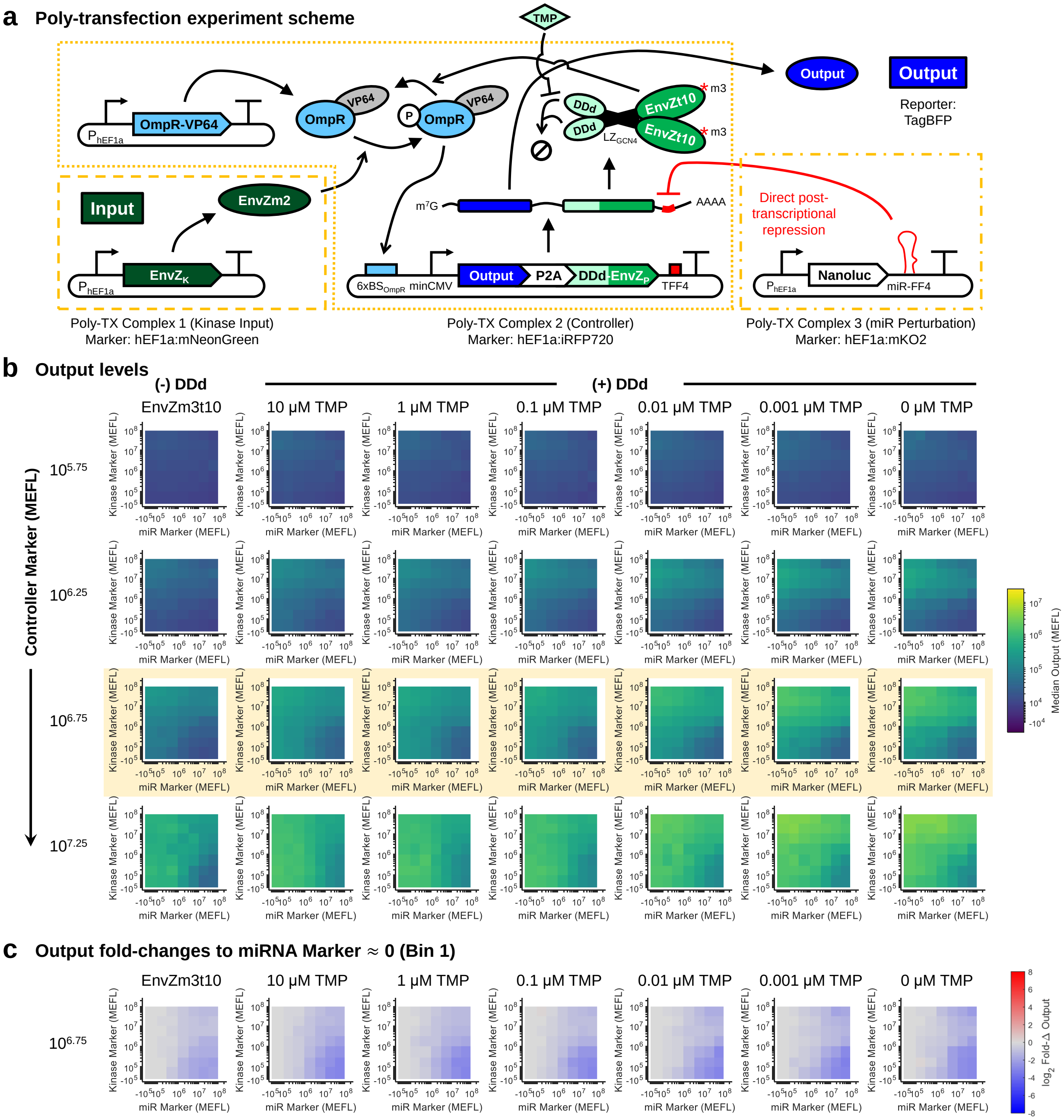


**Supplementary Figure 24: Extended analysis of CL variants with miR-FF4 perturbation**. **(a-c)** Similar analysis as in Supplementary Figure 23 for CL variants with DDd-EnvZm3t10 as the phosphatase (DDd-CL) and at different dosages of TMP. The data for the CL system without DDd is provided for reference. The highlighted plots in Panel (b) correspond to the data shown in Panel (c), Figures 27-5 & 6, and Supplementary Figures 29. All data was collected 48 hours post-transfection in HEK-293FT cells.

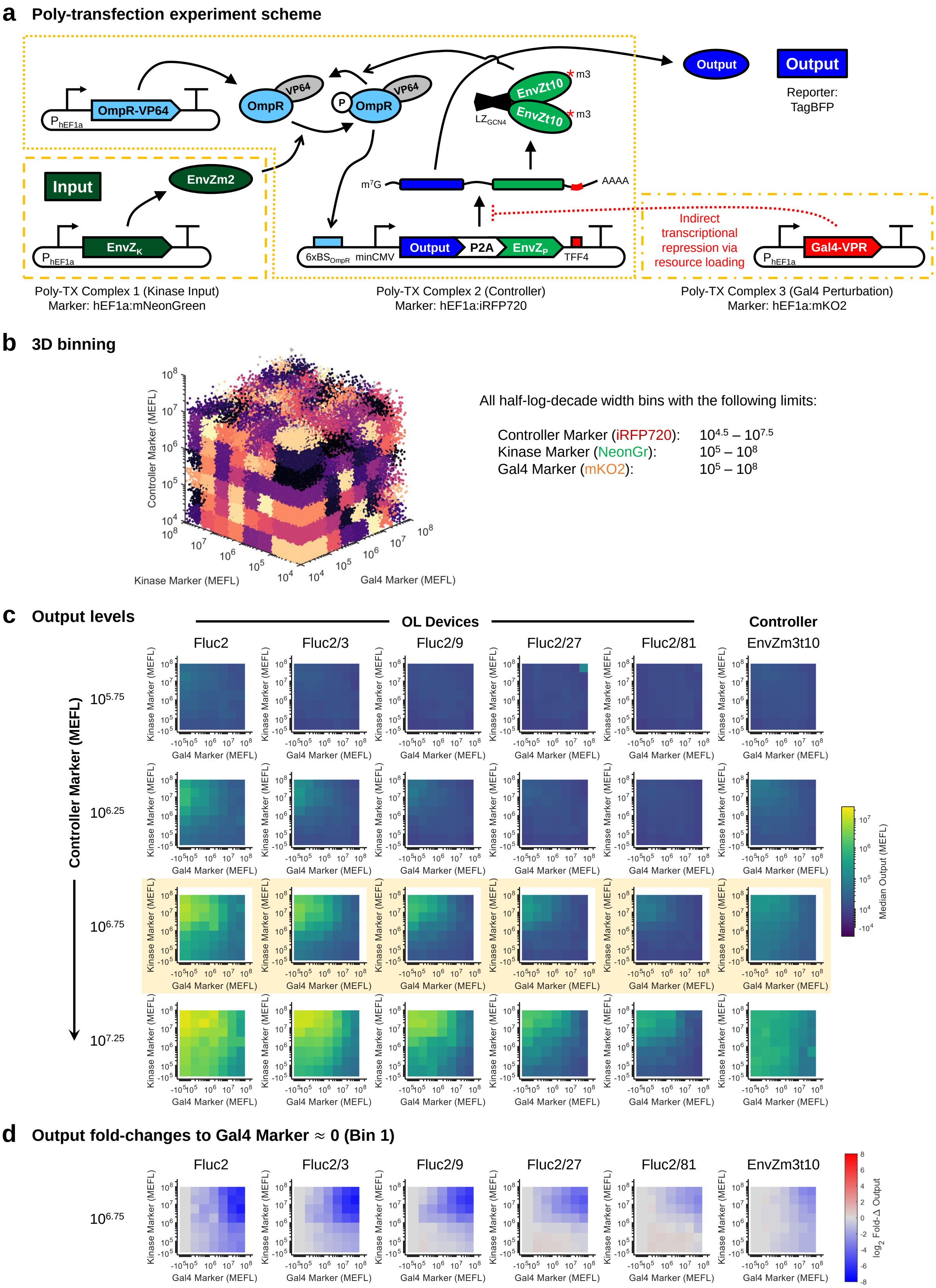


**Supplementary Figure 25: Extended analysis of OL variants with Gal4-VPR perturbation**

**Supplementary Figure 25:** (Previous page.) **(a-d)** Analogous plots to Supplementary Figure 23, but with Gal4-VPR perturbation rather than miR-FF4. Gal4-VPR knocks down output expression via loading of transcriptional resources[51]. The highlighted plots correspond to the data shown in Panel (d), Figures 5 & 6, and Supplementary Figures 27 & 30.

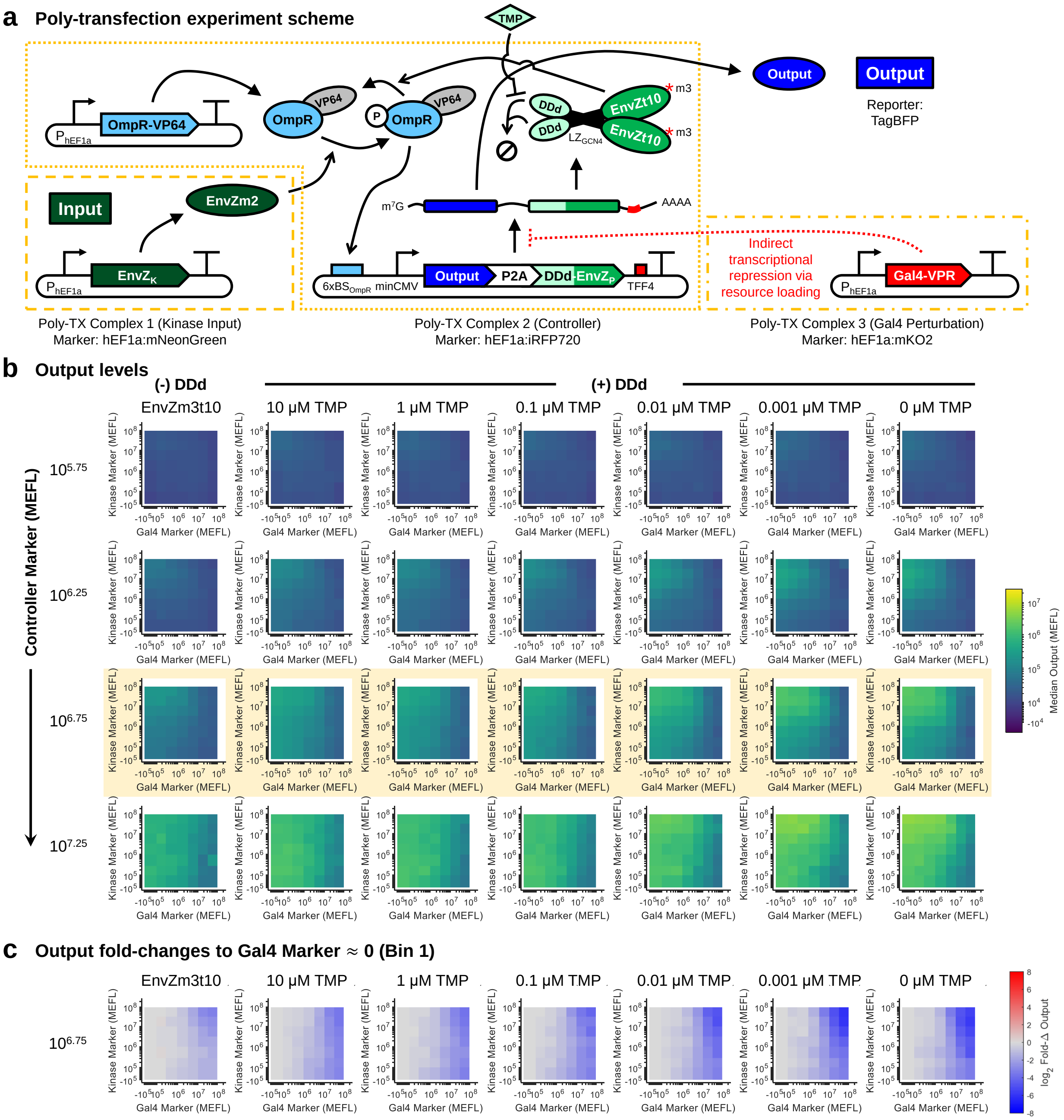


**Supplementary Figure 26: Extended analysis of CL variants with Gal4-VPR perturbation**. **(a-c)** Similar analysis as in Supplementary Figure 25, but with DDd-CL at different TMP concentrations. The data for the CL system without DDd is provided for reference. The highlighted plots in Panel (b) correspond to the data shown in Panel (c), Figures 5 & 6, and Supplementary Figures 27, 28, & 30. All data was collected 48 hours post-transfection in HEK-293FT cells.

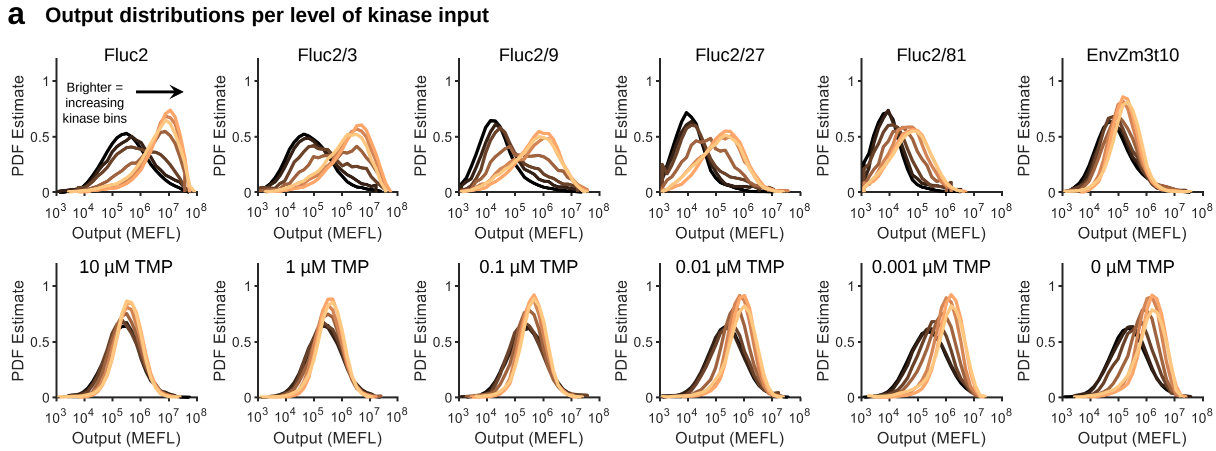


**Supplementary Figure 27: Distribution of output expression across kinase input levels**. **(a)** Distributions for the bins highlighted in Supplementary Figure 23c for the bins without miR-FF4 (miR Bin 1: 0 MEFLs). Brighter colored lines correspond to bins with increasing kinase. Data is representative from the first experimental repeat. Data in this figure is extracted from the highlighted bins in Supplementary Figures 23-26.

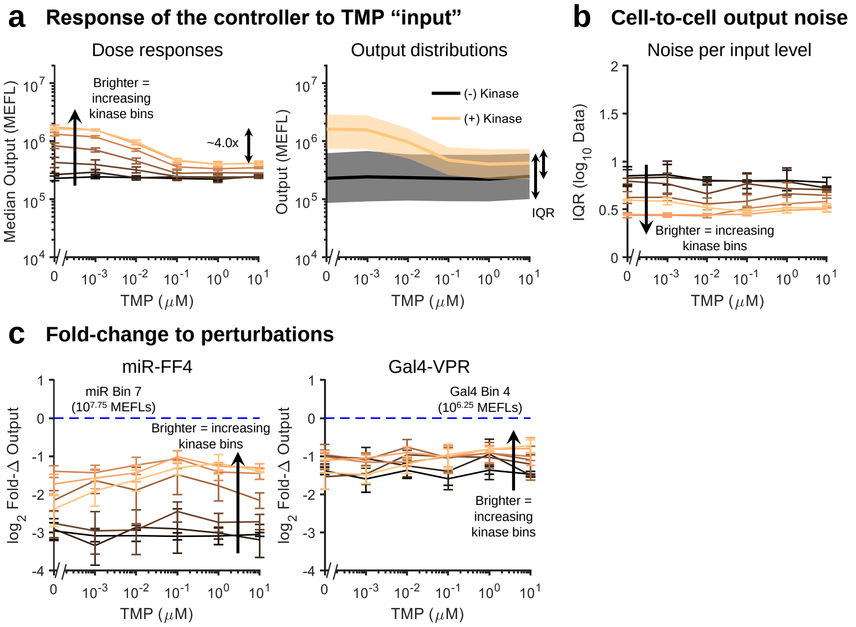


**Supplementary Figure 28: Response of DDd-CL system to TMP input**. **(a)** Dose-response of the DDd-CL system to TMP – comparable to Figure 5c. **(b)** Quantification of cell-to-cell noise in output levels of the DDd-CL system as a function of TMP dosage. **(c)** Fold-changes of the DDd-CL system to perturbations as a function of TMP dosage. Data in this figure is extracted from the highlighted bins in Supplementary Figures 24 & 26.

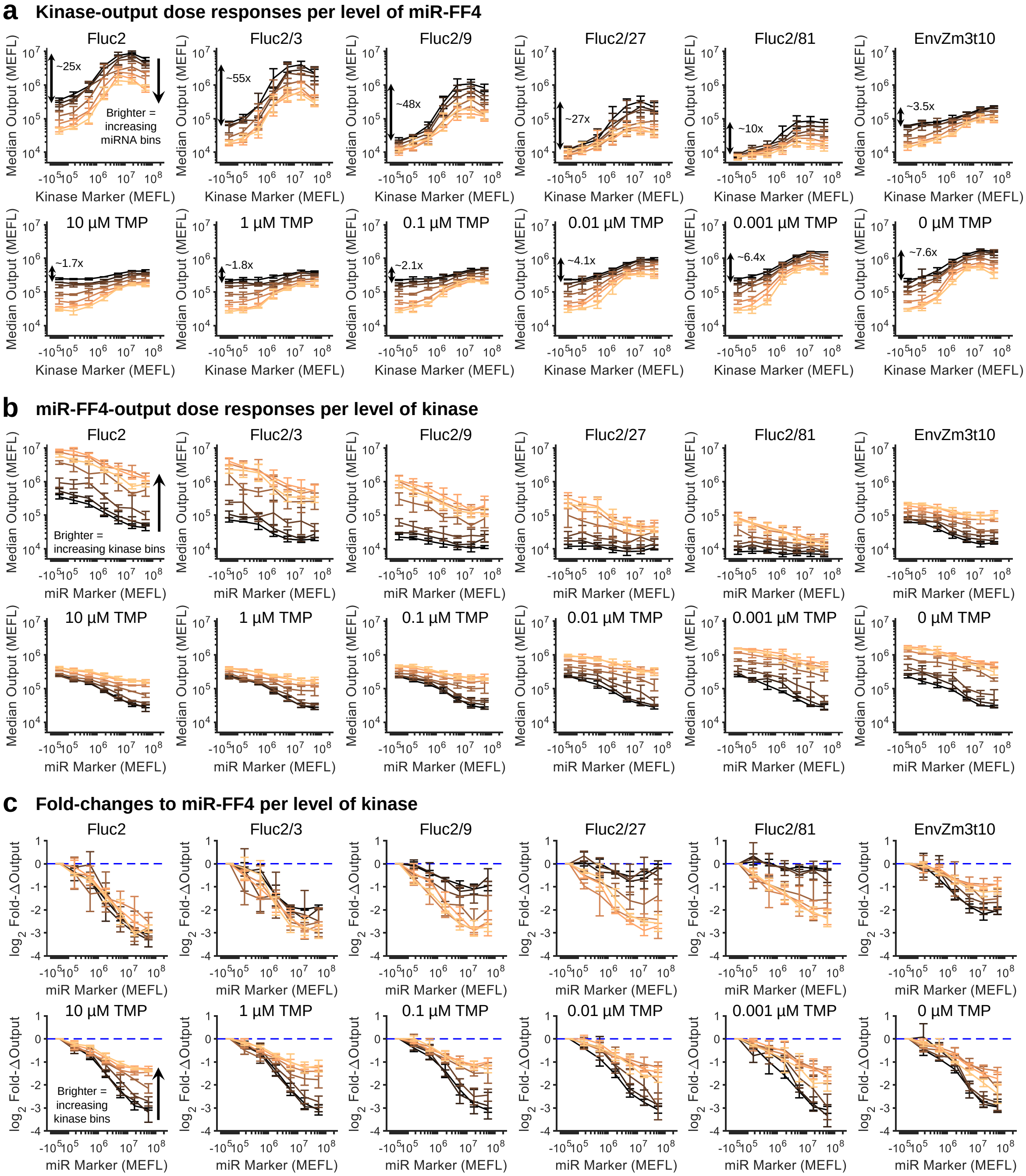


**Supplementary Figure 29: Detailed comparison of OL responses to miR-FF4 perturbation (a)** Kinase-output dose-response curves for OL and CL systems, corresponding to the bins highlighted in Supplementary Figures 23 24. The numbers indicate the maximum fold-change due to addition of the kinase for the bins without miR-FF4 (miR Bin 1: 0 MEFLs). Brighter colored lines correspond to bins with increasing miR-FF4. **(b)** Response of output expression to miR-FF4. Brighter colored lines correspond to bins with increasing kinase. **(c)** Fold-change of output expression to miR-FF4. Brighter colored lines correspond to bins with increasing kinase. The dashed blue lines provide a reference no change (ideal). In Panels (a) and (b), errorbars indicate the mean ± s.d. of measurements from three experimental repeats.

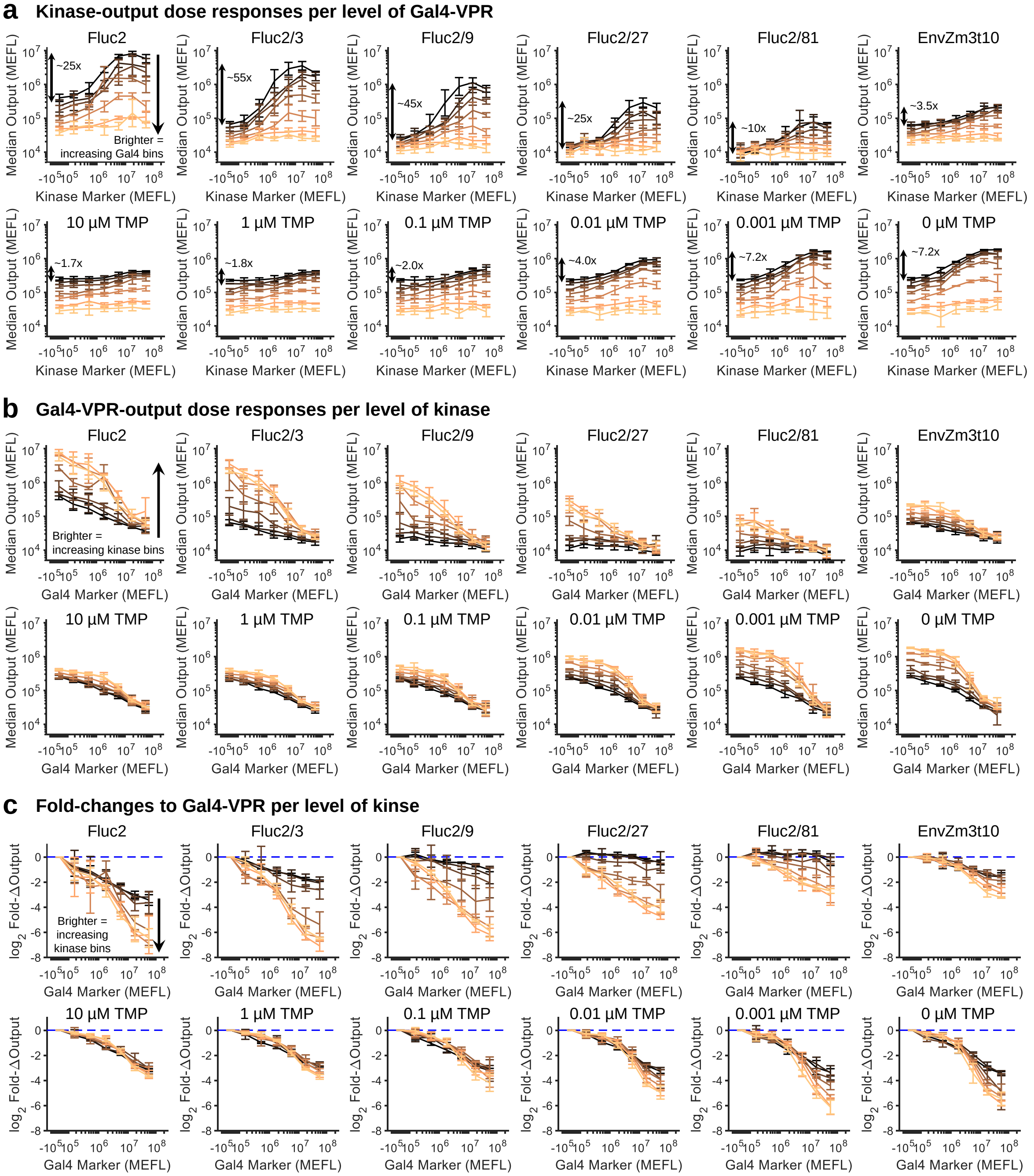


**Supplementary Figure 30: Detailed comparison of OL responses to Gal4-VPR perturbation (a-e)** Analogous plots to Supplementary Figure 29, but with Gal4-VPR perturbation rather than miR-FF4. The Gal4-VPR bin used for comparison was Bin 4 ($10^{6.25}$ MEFLs), which knocked down the OL samples to a similar degree as the highest bin for the miR-FF4 perturbation.

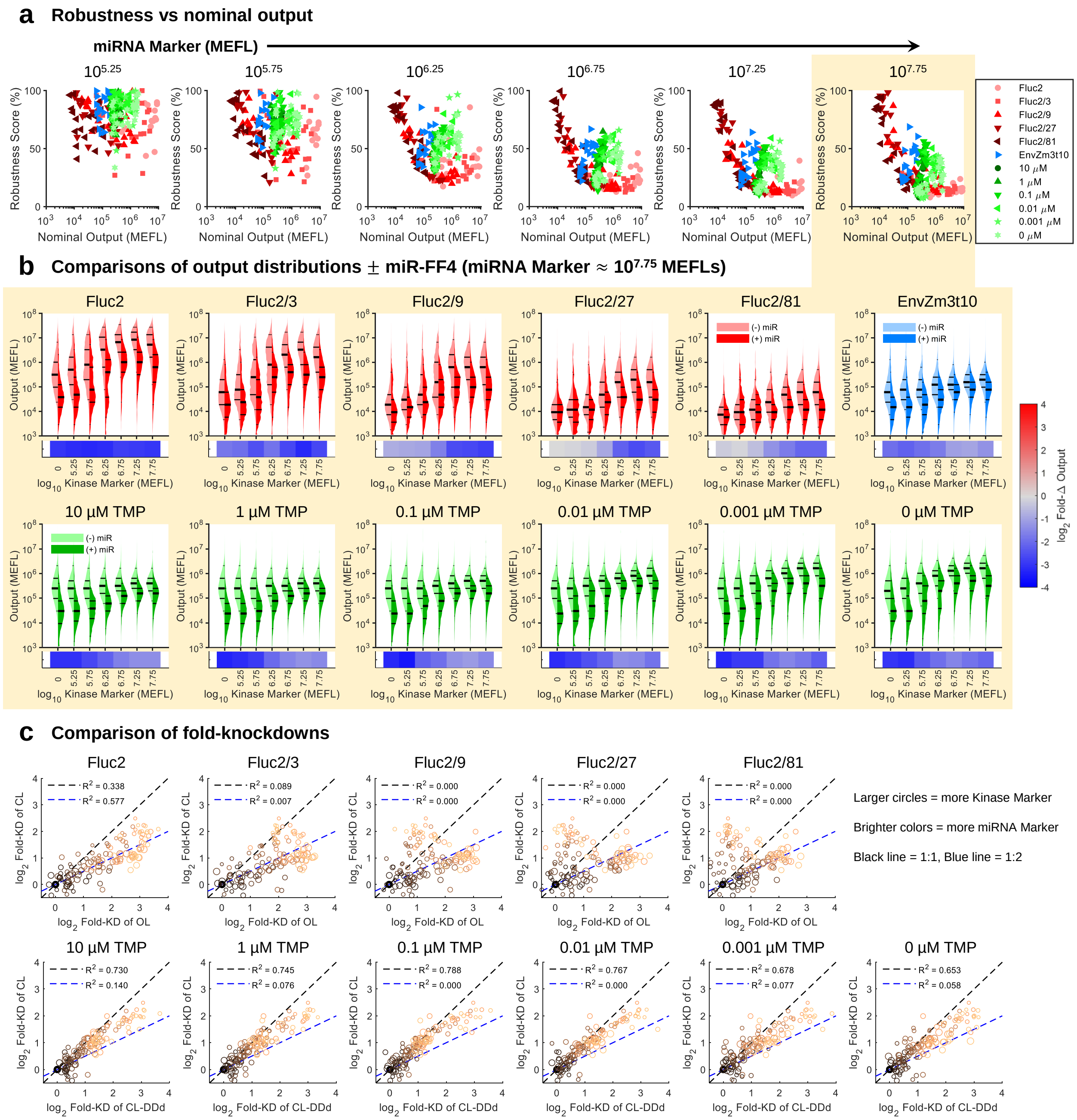


**Supplementary Figure 31: Detailed comparison of CL responses to miR-FF4 perturbation**

**Supplementary Figure 31:** (Previous page.) **(a)** Robustness scores (100% − % deviation due to perturbation) for all OL and CL systems, plotted separately for each non-zero miR-FF4 bin. The nominal output corresponds to the level of output in the absence of perturbation. **(b)** Comparison of output distributions per kinase bin in the absence (–, Bin 1) and presence (+, bin highlighted in (a)) of miR-FF4. Fold-changes are indicated in the heatmaps below the violin plots. The lines on the violins denote the $5^{th}$, $25^{th}$, $50^{th}$, $75^{th}$, and $95^{th}$ percentiles. **(c)** Comparison of fold-changes between each OL variant and the CL system without DDd (tow row), as well as between the CL systems with and without DDd (bottom row). Larger circles indicate more kinase and brighter colors indicate more miR-FF4. For levels of kinase that give output expression above background, the OL samples are knocked down approximately twice as much in log space, *i.e.* a squared fold-change in linear space. In Panels (a) and (c), points from each experimental repeat are plotted individually. In Panel (b), the data is representative of the first of three experimental repeat.

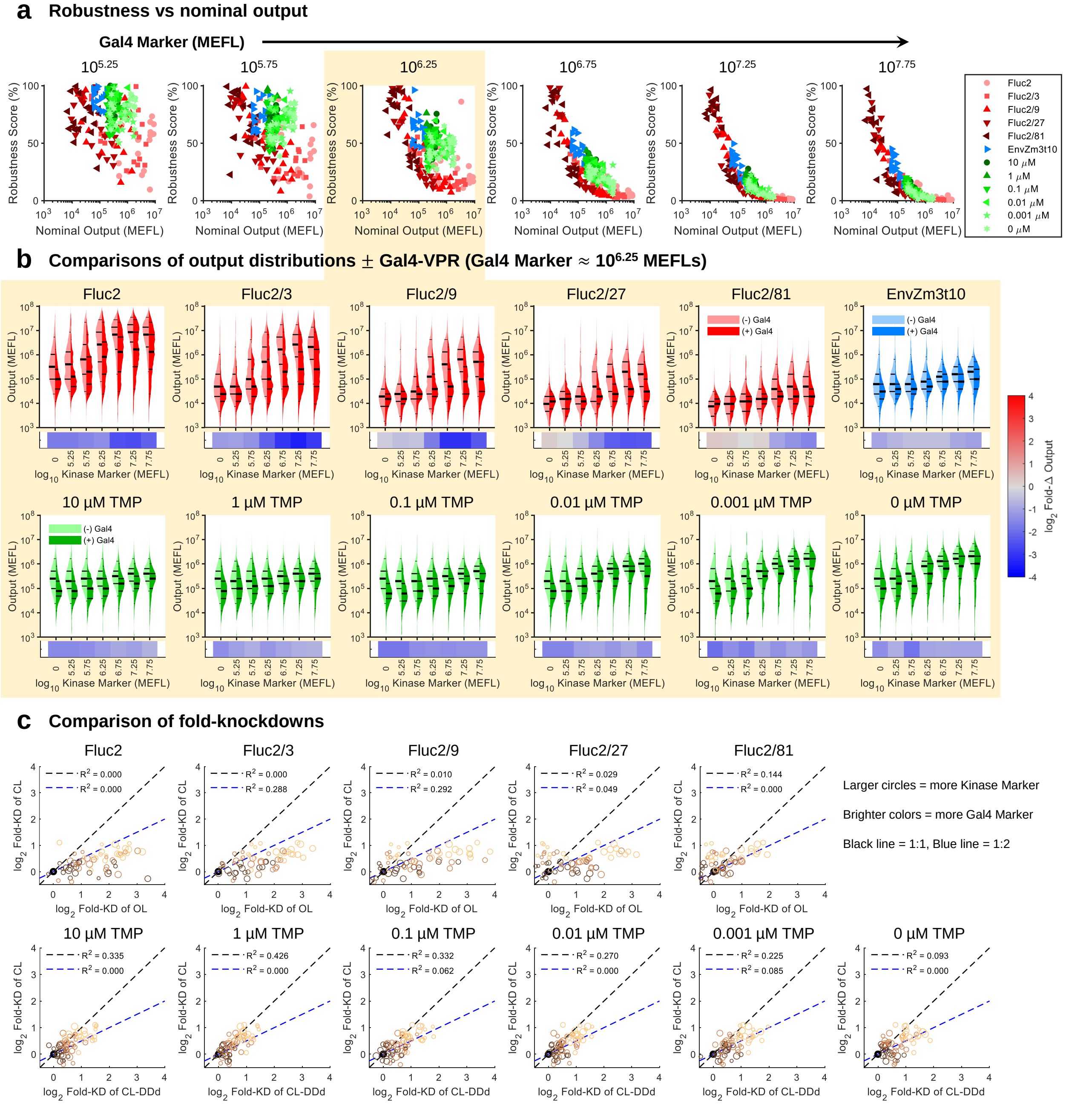


**Supplementary Figure 32: Detailed comparison of CL responses to Gal4-VPR perturbation (a-c)** Analogous plots to Supplementary Figure 31, but Gal4-VPR perturbation. Note that in Panel (c), we only compare bins of Gal4 up to that highlighted in the preceding panels.

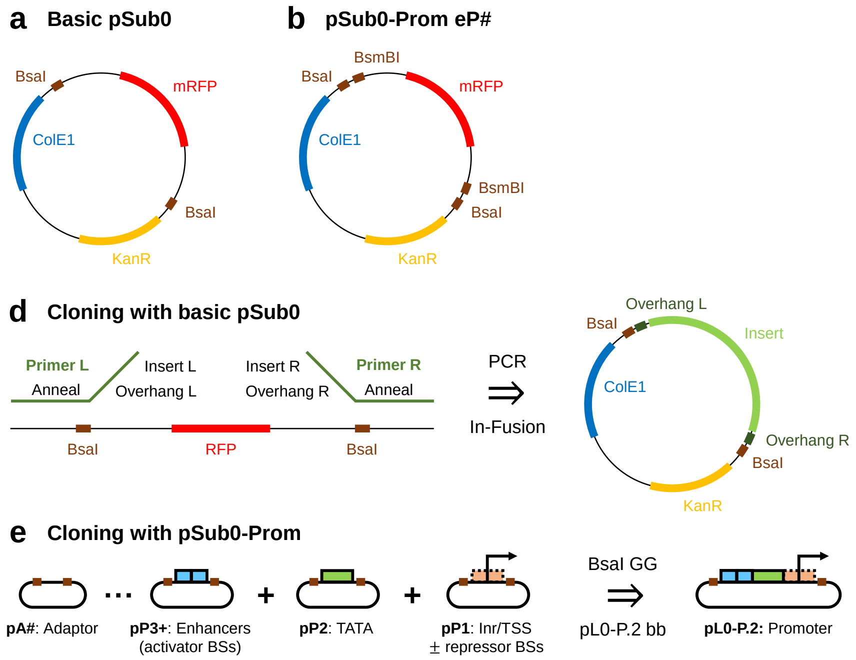


**Supplementary Figure 33: New pSub0 system for cloning promoters**. **(a-b)** Schematic of pSub0 backbones showing important cloning features. Note that the basic pSub0 (a) does not have 'real' overhangs. The BsmBI sites in the promoter (pSub0-Prom-pP# [b]) position vectors are for inserting TF binding sequences. Successfully-cloned pSub0 parts can be identified by red/white colony screening via replacement of mRFP. **(c)** Cloning with basic pSub0 is conducted by PCR of the pSub0 template; overhangs and the insert are added in via the 3' extension of the primers. Typically, PCR is followed with In-Fusion, which required just a 15 bp overlap of the insert by each primer extension. This method is used to create small pSub0s that do not require re-positioning, *e.g.* minimal promoter elements. **(d)** Cloning strategy to make promoters from pSub0s. Unlike typical Moclo, promoters are constructed from the 3' to 5' end, since the most conserved sequences are at the distal 3' end of the promoter. The pSub0s are cloned together into a promoter using BsaI Golden Gate, but can also be directly combined with other pL0s to make a pL1 (Since they have BsaI overhangs). Because the pL0 backbones are not optimal for insertion with BsaI Golden Gate, it is more efficient to make pL0s from pSub0s by first doing Golden Gate with the inserted parts, then ligating into a pre-fragmented pL0 backbone. Overhangs for all pSub0s are provided in Table 1.

**Supplementary Table 1: pSub0-Promoter overhangs**. Note that the sequences of many plasmids used in experiments have older versions of these overhangs. Note also that overhang 5 (between pP5 and pP6) is incompatible with pL0 → pL1 cloning, so pSub0-Prom parts up to pP4 can be cloned directly in place of a pL0 P.2 part.

| **Position** | **5'** | **3'** |
|---|---|---|
| pP1 | ACGC | CAGA |
| pP2 | TGTG | ACGC |
| pP3 | CTGA | TGTG |
| pP4 | TAAA | CTGA |
| pP5 | AGGT | TAAA |
| pP6 | AAGA | AGGT |
| pP7 | ACCG | AAGA |
| pP8 | TAGA | ACCG |
| pP9 | CTAA | TAGA |
| pP10 | CCCT | CTAA |
| pA# | TACT | NNNN |
